# BMAL1 represses transposable elements independently of CLOCK in pluripotent cells

**DOI:** 10.1101/2024.06.18.599568

**Authors:** Amador Gallardo, Efres Belmonte-Reche, María Marti-Marimon, Joan Domingo-Reinés, Guillermo Peris, Lourdes López-Onieva, Iván Fernández-Rengel, Pablo Tristán-Ramos, Nicolas Bellora, Antonio Sánchez-Pozo, Antonio M Estévez, Sara R Heras, Marc A. Marti-Renom, David Landeira

## Abstract

Circadian oscillations of gene transcripts rely on a negative feedback loop executed by the activating BMAL1-CLOCK heterodimer and its negative regulators PER and CRY. Although circadian rhythms and CLOCK protein are mostly absent during embryogenesis, the lack of BMAL1 during prenatal development causes an early aging phenotype during adulthood, suggesting that BMAL1 carries out an unknown non-circadian function during organism development that is fundamental for healthy adult life. Here, we show that BMAL1 interacts with TRIM28 and represses transcription of totipotency-associated MERVL retrotransposons in mouse pluripotent cells. Deletion of Bmal1 leads to genome-wide upregulation of MERVLs, changes in the three-dimensional organization of the genome, and acquisition of totipotency-associated features. Overall, we demonstrate that in pluripotent cells BMAL1 is redeployed as a transcriptional repressor of transposable elements (TEs) in a CLOCK-independent way. We propose that BMAL1-TRIM28 activity during prenatal life is essential for optimal health and life span in mammals.

## Introduction

Organisms that live on the surface of the earth are regulated by circadian rhythms that optimize their physiology during daily changes in sunlight ^1^. In mammals, this is achieved through specialized photoreceptor cells in the retina that transmit information about light intensity to the brain, which then synchronizes peripheral molecular clocks present is most cells of the adult organism through humoral signals ^1, 2^. The molecular clock is based on the core heterodimeric protein BMAL1-CLOCK that activates transcription of its own negative regulators PER1/2 and CRY1/2 genes, generating a negative feedback loop that produces daily oscillations in the expression of up to 20 % of cellular transcripts in a tissue-specific way (also known as clock-controlled genes, CCGs) ^3, 4^. Dysregulation of CCGs during adulthood is commonly assumed to underlie the wide range of health disorders associated to misfunctioning of the circadian machinery ^5, 6, 7^. However, the phenotypes of mice mutant for Clock and Bmal1 genes are ostensibly dissimilar - altered metabolism and obesity in Clock mutant mice ^8^ versus early aging phenotype in Bmal1^-/-^ specimens ^9^. The differences observed in these phenotypes indicate that many pathological conditions associated to the molecular clock might be a consequence of mutually independent functions of BMAL1 and CLOCK. Moreover, differences in the phenotypes of Clock and Bmal1 mutant mice emerge from the activity of BMAL1 during embryo development, because most of the phenotype observed in Bmal1^-/-^ adult mice ^9^ is caused by the absence of BMAL1 during prenatal development, but not during adulthood ^10^. Importantly, the molecular mechanism by which BMAL1 influences mouse development is unknown, but it is likely independent of its canonical function as an oscillator of gene transcription, because robust circadian oscillations of gene transcripts are not detected until perinatal stages when CLOCK protein becomes robustly expressed^11, 12, 13, 14, 15, 16, 17, 18^.

To investigate the alternative non-circadian function of BMAL1 during embryogenesis we focused our analysis on early stages of mouse development. Following fertilization, the mouse zygote divides, and its genome is activated producing two totipotent cells (2C) that differentiate into embryonic or extraembryonic tissues ^19^. The totipotent state is characterized by the expression of 2C-specific genes and Murine Endogenous Retroviral Element with a Leucine tRNA primer binding site retrotransposons (MuERV-L; also known as MERVLs), that are transcriptionally repressed when cells differentiate into either trophectoderm or the pluripotent inner cell mas (ICM) that form the early blastocyst ^20, 21^. The molecular basis of totipotency is poorly understood, due to difficulties in the isolation of totipotent cells directly from the 2C stage blastomeres ^21^. However, cultures of pluripotent embryonic stem cells (mESCs) can be easily derived from the ICM of the blastocyst ^22^, and they contain a fraction of cells in a metastable state that display molecular features that resemble the totipotent 2C blastomeres (2C- like cells (2CLC)) ^21, 23^. Thus, mESCs cultures are an amenable in-vitro system to dissect the molecular basis of totipotency to pluripotency transition.

Here, we have examined the function of BMAL1 in pluripotent mESCs. We show that BMAL1 interacts with TRIM28, facilitates H3K9me3 deposition and represses the transcription of transposable elements (TEs) in mESCs. Importantly, deletion of Bmal1 leads to the transcriptional activation of MERVL copies, changes in the three-dimensional (3D) organization of the genome and acquisition of 2C-associated cellular and molecular features. Overall, we found that in the absence of CLOCK protein BMAL1 is redeployed as a transcriptional repressor that regulates the expression of TEs and 3D genome organization in pluripotent cells. We propose that unintended expression of retrotransposons in Bmal1^-/-^ mice alters the formation of healthy tissue during development, and that this is later reflected as premature aging during adulthood. Our discoveries establish a novel molecular framework for future studies dissecting the molecular details by which the lack of BMAL1 during embryogenesis results in accelerated aging later during adulthood.

## Results

### BMAL1 is required for the recruitment of TRIM28 to chromatin

Analysis of our published mRNA sequencing (mRNA-seq) datasets of parental (JM8^+/+^) and Bmal1^-/-^ mESCs ^16^ revealed that mis-expressed genes in Bmal1^-/-^ cells are over-expressed (454 upregulated vs 116 downregulated, FC>2, p<0.05) (Figure 1A, Table S1), supporting that BMAL1 is not acting as a transcriptional activator in partnership with CLOCK during early development. To address the molecular mechanism by which BMAL1 represses gene transcription in mESCs we established a clonal cell line that expresses a FLAG-tagged version of BMAL1 (FLAG-BMAL1), isolated nuclei, and analysed the interactome of BMAL1 by anti-FLAG co-immunoprecipitation followed by mass spectrometry (Figure S1A – S1C). We found that FLAG-BMAL1 interacts with factors involved in transcriptional repression (Table S2) such as TRIM28 (also known as KAP1) ^24^, NACC1 ^25^ and RIF1 ^26^, and several zinc finger proteins (ZFPs) including ZFP638, ZFP326 and ZFP281 (Figure S1D). We also identified interactions with proteins involved in rRNA production and mRNA splicing (Table S2), but these were not further pursued because they are common contaminants in this type of purification schemes and neither rRNA expression nor mRNA splicing was affected in Bmal1^-/-^ cells (Figure S1E – S1G).

**Figure 1.**
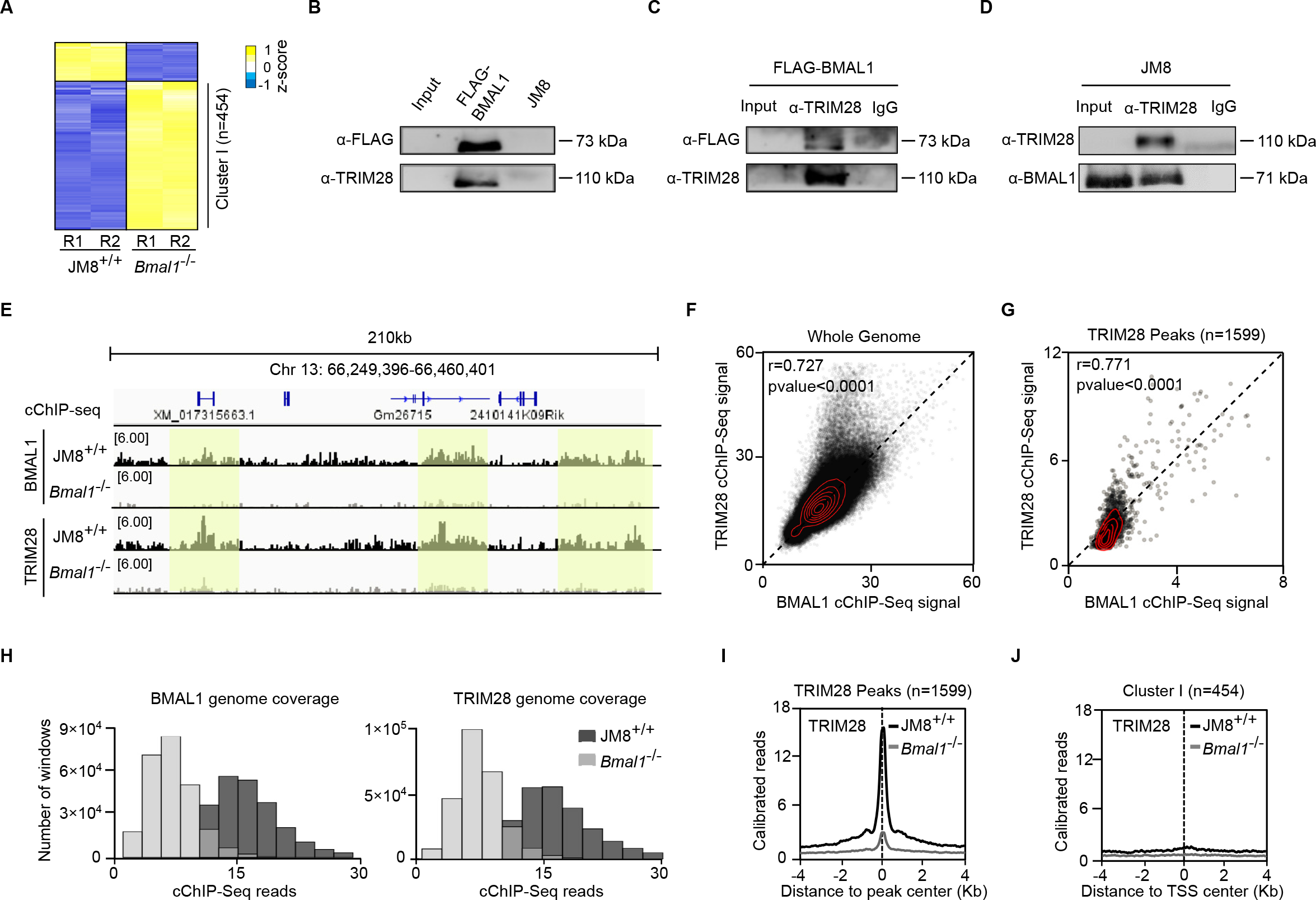
BMAL1 is required for the recruitment of TRIM28 to chromatin (A) Heatmap of genes mis-expressed in Bmal1^-/-^ compared to wild-type JM8^+/+^ mESCs analysed by mRNA-seq (fold change > 2 and p-value < 0.05). Yellow and blue colours indicate the Z-score (deviation relative to the average value). R1 and R2 indicate biological replicates of the indicated sample. (B) Co-Immunoprecipitation analysis of the interaction between BMAL1 and TRIM28 proteins in FLAG- BMAL1-expressing mESCs. An anti-FLAG antibody was used to pull down FLAG-BMAL1 and the interaction with TRIM28 was tested by western-blot. An immunoprecipitation using the anti-FLAG antibody in JM8 mESCs served as a negative control. (C) As in B but TRIM28 was immunoprecipitated using an anti-TRIM28 antibody and the interaction with FLAG-BMAL1 was analysed using an anti-FLAG antibody. Immunoprecipitation with an anti-IgG antibody was used as a negative control. (D) Co-immunoprecipitation analysis by western blot of TRIM28 and endogenous BMAL1 in JM8 mESCs. Anti-IgG immunoprecipitation was used as negative control. (E) Genome browser view of BMAL1 and TRIM28 binding profiles in JM8^+/+^ and Bmal1 ^-/-^ mESCs. Light yellow rectangles highlight the genomic locations with punctate high-level enrichment of BMAL1 and TRIM28 binding. (F) Plot showing the genome-wide correlation of TRIM28 and BMAL1 cChIP-seq signals in mESCs. The genome was divided in adjacent 10kb-sized windows for which total reads were calculated and plotted. Pearsońs correlation coefficient (r) and p-value are indicated. (G) Correlation plot of BMAL1 and TRIM28 cChIP-seq signals at all the regions identified as peaks (n=1599) in the TRIM28 cChIP-seq. Pearsońs correlation coefficient (r) and p-value are indicated. (H) Histogram showing the genome-wide level of BMAL1 and TRIM28 in JM8^+/+^ and Bmal1^-/-^ mESCs. Number of windows for a given total read signal upon dividing the genome in 10kb-sized adjacent windows is plotted. (I) Average binding profiles of TRIM28 in JM8^+/+^ and Bmal1^-/-^ mESCs around the centre of TRIM28 binding peaks (n=1599) identified in JM8^+/+^ cells. (J) Average binding profiles of TRIM28 in JM8^+/+^ and Bmal1^-/-^ mESCs around the TSS of genes that are upregulated in Bmal1^-/-^ mESCs (n=454).

Among transcriptional repressors identified (Figure S1D), we focused on TRIM28 because it has a well- stablished role acting as a scaffold protein that recruits the chromatin regulators NuRD and SETDB1 to genomic regions ^24, 27, 28, 29, 30^. As expected, protein coimmunoprecipitation followed by western blot confirmed that FLAG-BMAL1 and endogenous BMAL1 interacts with TRIM28 (Figure 1B-1D). To address whether BMAL1 and TRIM28 bind to the same regions of the genome, and whether they are functionally linked, we performed calibrated chromatin immunoprecipitation followed by sequencing (cChIP-seq) using antibodies against BMAL1 or TRIM28 in JM8^+/+^ parental wild-type and Bmal1^-/-^ mESCs. Inspection of cChIP-seq signals in the genome browser revealed that BMAL1 and TRIM28 display a visually concordant distribution across the genome, although the accumulation of BMAL1 at punctate high-level target regions is less obvious when compared to the signal observed for TRIM28 (Figure 1E). Importantly, low-level ubiquitous binding of BMAL1 and TRIM28 is evident in JM8^+/+^ but not in Bmal1^-/-^ cells, demonstrating that BMAL1-TRIM28 complexes bind to DNA sequences that are not marked by a punctate high-level signal (Figure 1E). Genome-wide binding analyses showed a positive correlation between BMAL1 and TRIM28 binding (Figure 1F and 1G), supporting that BMAL1 and TRIM28 have a largely overlapping distribution genome-wide. As expected, there was also a clear positive correlation in the binding signals of FLAG-BMAL1, BMAL1 and TRIM28 around the genome (Figure S1H and S1I). Importantly, the genome-wide binding signal of TRIM28 was reduced in Bmal1^-/-^ mESCs (Figure 1H-1I), indicating that BMAL1 is required to recruit TRIM28 to target chromatin. Of note, genes upregulated in Bmal1^-/-^ mESCs (Figure 1A) do not display a high-level signal of TRIM28 binding around their transcriptional start site (TSS) (Figure 1J), suggesting that BMAL1-TRIM28 complexes repress the transcription of protein coding genes through a mechanism that does not require their accumulation at gene promoters.

### BMAL1-TRIM28 modulates H3K9me3 levels and represses MERVL elements in mESCs

TRIM28 represses transcription of TEs in mESCs ^31, 32^, and TEs can impact the regulation of protein-coding genes through several mechanisms ^33^. Thus, we hypothesized that BMAL1-TRIM28 regulates the expression of TEs and indirectly modulate gene expression in mESCs. Peak calling analyses showed that regions with punctate high-level enrichment of TRIM28 (usually refereed as binding peaks) mainly fell into repeated DNA sequences (1378 out of 1599), with a strong bias (1333 out of 1378) towards retrotransposons that contain long terminal repeats (LTRs) (Figure 2A). Importantly, depletion of BMAL1 reduces the binding of TRIM28 to DNA repeats (Figure 2B) and promotes a drastic transcriptional upregulation of TEs copies genome-wide (1867 copies, FC>2, p<0.05) (Figure 2C) (Table S1), of which most (1399 out of 1867) were LTR retrotransposons (Figure 2D). Importantly, full-length MERVL elements were particularly overrepresented among the over-expressed TEs (605 copies) (Figure 2E and S2A). Notably, despite low expression of MERVLs makes them difficult to be detected in standard RNA-seq experiments in mESCs, we could readily detect upregulation of 24,2% of all annotated MERVL copies (605 of 2500) in Bmal1^-/-^ cells, suggesting that BMAL1 is involved in transcriptional repression of MERVL elements genome-wide. In agreement, complementary family-based analysis by mRNA-seq (Figure S2B and S2C) and RT-qPCR (Figure S2D) confirmed that BMAL1 is required to maintain MERVL elements transcriptionally repressed in mESCs.

**Figure 2.**
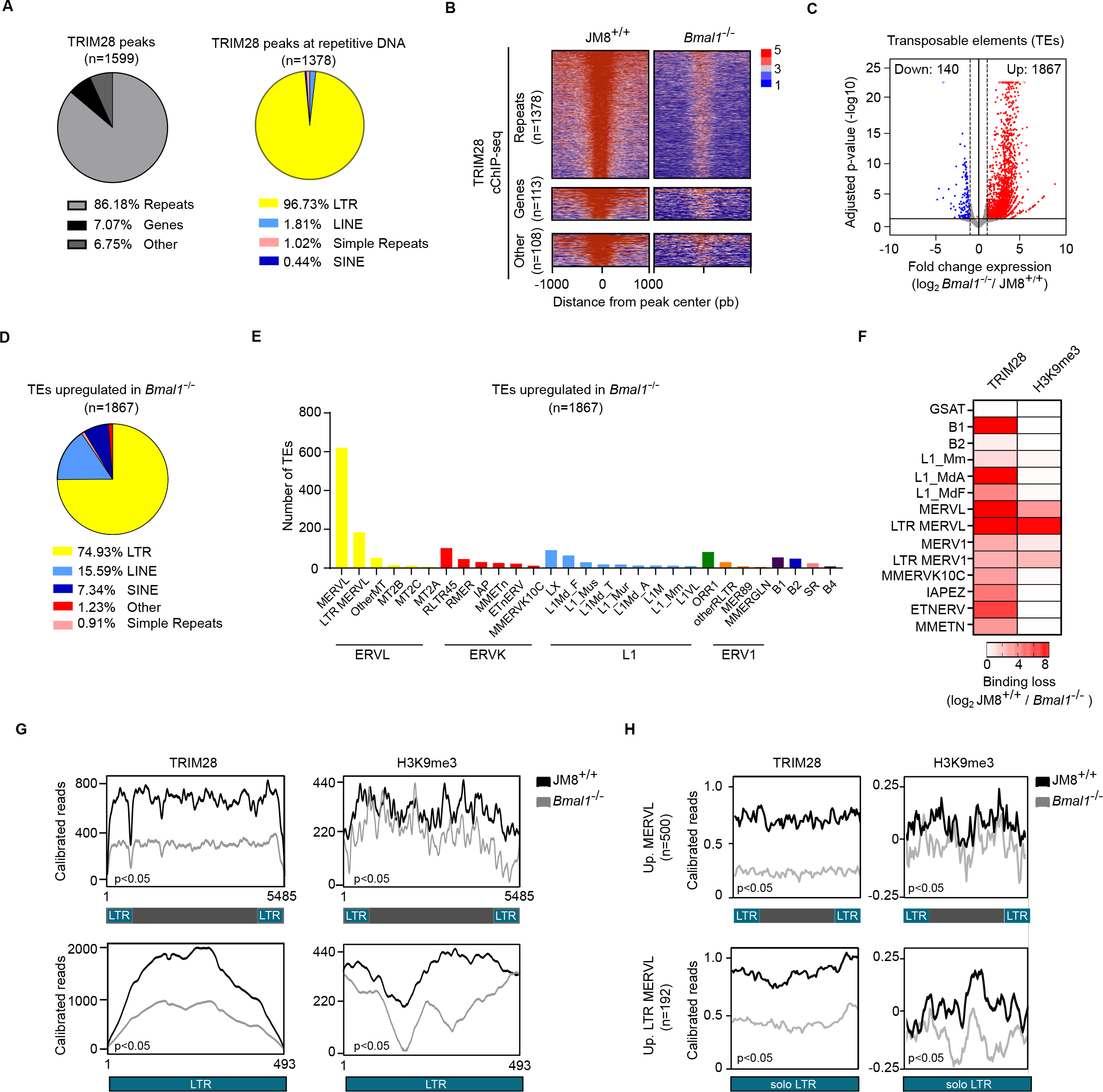
BMAL1 facilitates binding of TRIM28, H3K9me3 labelling, and transcriptional repression of MERVL elements in mESCs (A) Genomic distribution of TRIM28 binding peaks. (B) Heatmap comparing TRIM28 binding signal in JM8^+/+^ and Bmal1^-/-^ mESCs around the centre of the regions bound by TRIM28 in JM8^+/+^ cells and classified in indicated categories. (C) Volcano plot showing the expression level of individual TEs in Bmal1^-/-^ relative to JM8^+/+^ mESCs. Significant upregulation (FC>2, p-value<0.05, red colour), downregulation (FC<2, p-value<0.05, blue colour) or no change is indicated (grey colour). (D, E) Pie diagram (D) and bar plot (E) showing the classification of the 1867 TEs that are overexpressed in Bmal1^-/-^ mESCs. (F) Heatmap showing the reduction of TRIM28 binding and H3K9me3 in Bmal1^-/-^ mESCs (relative to JM8^+/+^), as assayed by cChIP-seq, at the consensus sequence of indicated families of TEs. (G) Plots displaying the TRIM28 and H3K9me3 enrichment signals along the consensus sequence of full MERVL (top panels) or MERVL-specific LTR (also known as MT2_mm) elements (bottom panels) in JM8^+/+^ and Bmal1^-/-^ mESCs. p value was calculated using an ANOVA test. (H) Plots displaying the TRIM28 and H3K9me3 enrichment profile in JM8^+/+^ and Bmal1^-/-^ cells along the MERVL (top panels) or solo MERVL-specific LTR (also known as MT2_mm) (bottom panels) that were upregulated in Bmal1^-/-^ mESCs. p value was calculated using an ANOVA test.

TRIM28 interacts with SETDB1 and facilitates methylation of H3K9 and transcriptional repression of endogenous retroviruses (ERVs) in mESCs ^28, 31, 32^. Thus, we hypothesized that in the absence BMAL1 and efficient TRIM28 recruitment (Figure 1), the distribution of H3K9me3 would be perturbed and facilitate transcriptional upregulation of MERVL elements. Global levels of TRIM28 protein and H3K9me3 remain mostly unaltered in BMAL1-depleted cells (Figure S2E). However, their accumulation within the nucleus (Figure S2F and S2G) and their presence on chromatin genome-wide (Figure 1H and S2H) was reduced in Bmal1 ^-/-^ cells compared to JM8^+/+^ cells. Notably, in the absence of BMAL1, binding of TRIM28 was reduced at several families of TEs, but H3K9me3 was drastically reduced distinctively at MERVL elements (Figure 2F), indicating that BMAL1 is critical for maintaining TRIM28 binding and H3K9me3 levels mostly at MERVL elements. In keeping, enrichment analyses of TRIM28 binding and H3K9me3 decoration along MERVL and solo LTR sequences (consensus and over-expressed ones) confirmed their reduction in Bmal1^-/-^ cells compared to JM8^+/+^ cells (Figure 2G and 2H). Expression of H3K9 methyltransferases and demethylases was unaltered in Bmal1^-/-^ cells (Figure S2I), discarding that deregulation of MERVLs in Bmal1^-/-^ cells is an indirect effect of transcriptional mis-expression of genes involved in H3K9 methylation. Thus, we concluded that BMAL1 is required for efficient recruitment of TRIM28, maintenance of H3K9me3 levels and preservation of the transcriptionally repressed state of MERVL retrotransposons in mESCs.

### Transcriptional activation of MERVLs and neighbouring genes is associated to changes in 3D chromatin organization in Bmal1^-/-^ mESCs

Genes upregulated in Bmal1^-/-^ cells (Figure 1A) tend to be in linear proximity of upregulated MERVLs (median distance 121 Kb) as compared to unaltered control genes (median distance 637 Kb) (Figure 3A and 3B), supporting that production of RNA coming from MERVL and nearby genes is co-regulated in cis. Based on previous literature, we hypothesized that co-regulation of genes and MERVL elements could be based on the production of MERVL-containing chimeric transcripts ^23, 34^ or be associated to changes in 3D chromatin organization ^35, 36^.

**Figure 3.**
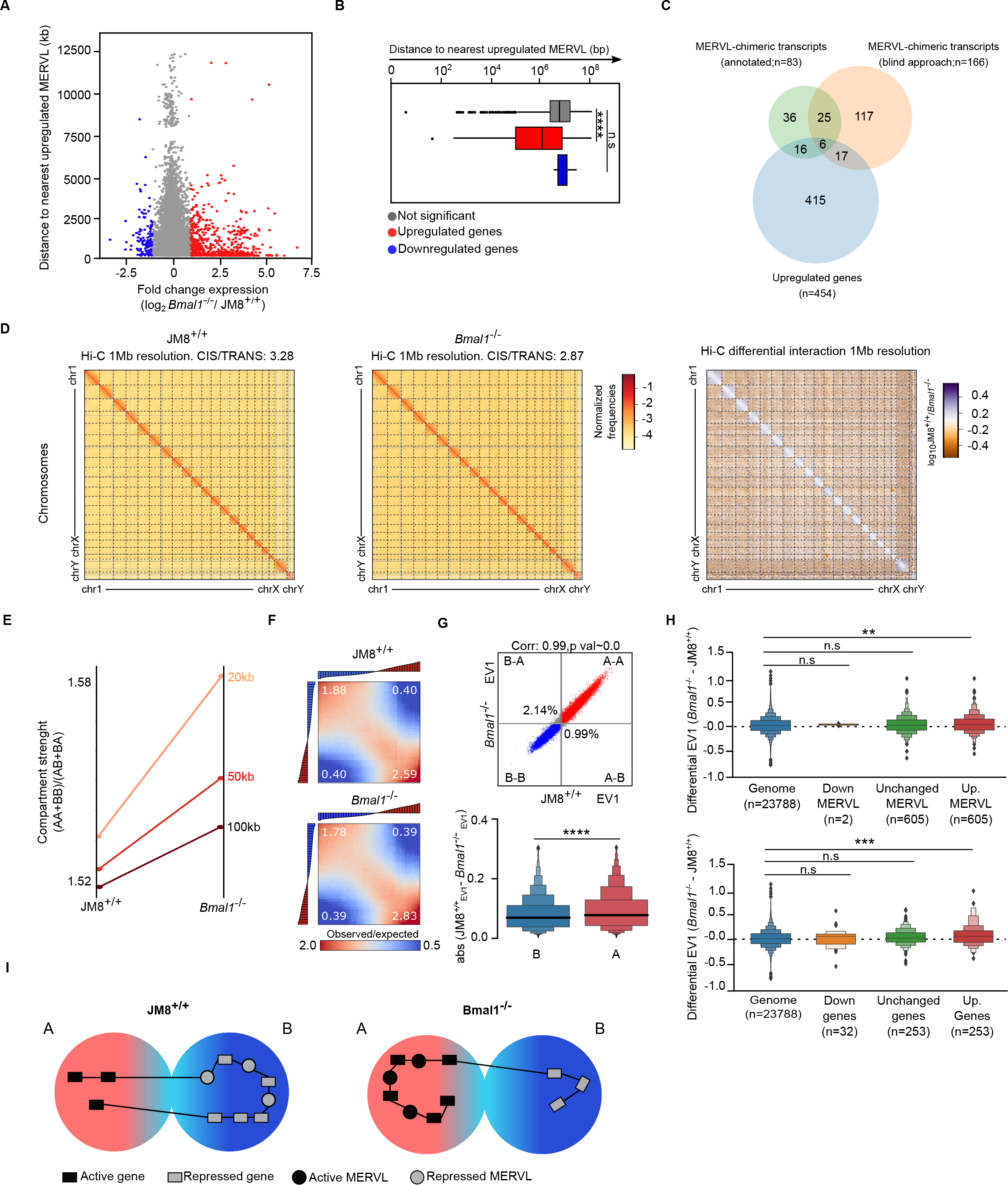
Transcriptional activation of MERVL retrotransposons and neighbouring genes is associated to changes in 3D chromatin organization in Bmal1^-/-^ mESCs (A) Plot showing the expression of protein coding genes in Bmal1^-/-^ cells (relative to JM8^+/+^) (x-axis) against their distance to the nearest upregulated MERVL in Bmal1^-/-^ mESCs. Upregulated (FC>2, p<0.05) or downregulated (FC<2, p<0.05) genes are marked as red or blue dots respectively. (B) Box plot comparing the distance between mis-expressed genes and the nearest upregulated MERVL in Bmal1^-/-^ cells. Genes upregulated (FC>2, p<0.05), downregulated (FC<2, p<0.05), or unchanged are shown. (C) Venn diagram showing the overlap between expressed MERVL-containing chimeric RNA transcripts and transcriptionally upregulated genes in Bmal1^-/-^ mESCs. (D) Genome-wide Hi-C interaction maps at 1Mb resolution for JM8^+/+^ and Bmal1^-/-^ cells (left and middle panels). Right panel shows the differential interaction (ratio) map between both signals. (E) Plot comparing the compartment strength at indicated resolutions in JM8^+/+^ and Bmal1^-/-^ cells. (F) Saddle plots depicting the level of contact interactions between A (red) and B (blue) compartments, as organized by eigenvector ranking, in JM8^+/+^ and Bmal1^-/-^ cells. (G) Correlation plot showing eigenvector values in JM8^+/+^ and Bmal1^-/-^ cells for genomic bins that stay in A compartment (red), B compartment (blue) or change (grey) (top panel). The percentage of changing compartments are indicated. Bottom panel displays the distribution of the difference in the eigenvector value (absolute value of JM8^+/+^ minus Bmal1^-/-^ signal) of bins categorized as A (red) or B (blue) in Bmal1^-/-^ cells. (H) Top panel shows a plot measuring compartment changes as the difference in the eigenvector value for bins that include all the genome, downregulated (log2FC<-1, p<0.05), upregulated (log2FC>1, p<0.05) or unchanged MERVL elements in Bmal1^-/-^ cells. Bottom panel shows a plot measuring compartment changes as the difference in the eigenvector value for bins that include all the genome, downregulated (log2FC<-1.5, p<0.05), upregulated (log2FC>1.5, p<0.05) or unchanged protein coding genes in Bmal1^-/-^ cells (I) Model proposing how the lack of BMAL1 generates a coordinated shift in the 3D distribution of MERVLS and protein coding genes towards A compartments where their expression is transcriptionally induced. In B, G and H asterisk represents statistically significance by a Mann Whitney test. ** p value <0.01, *** p value <0.001 and **** p value <0.0001.

To test the former hypothesis, we identified TEs-chimeric RNAs produced in Bmal1^-/-^ mESCs and determined that MERVL containing chimeric RNAs expressed in Bmal1^-/-^ did not match the sequences of protein coding genes upregulated in Bmal1^-/-^ cells (Figure 3C). In addition, distance- and strand-specific analysis of the closest protein coding gene TSS located upstream or downstream of upregulated MERVLs (605 MERVL copies with FC>2, p<0.05) supported that the 454 genes upregulated in Bmal1^-/-^ mESCs are not the result of chimeric transcription, because they are not located in linearly adjacent positions to upregulated MERVLs (Figure S3A, S3B and S3C). In keeping, MERVLs and solo LTRs that are transcriptionally-induced in Bmal1^-/-^ mESCs could be classified mostly as self-dependent, because they are encoded in intergenic regions or within gene bodies of protein coding genes that were not transcriptionally activated in Bmal1^-/-^ mESCs (Figure S3D). Taken together, we concluded that the in cis co-regulation of MERVLs and protein coding genes in Bmal1^-/-^ cells is not mediated by the synthesis of MERVL-containing chimeric transcripts.

To study whether 3D genome organization underlies the co-regulation of MERVLs and protein coding genes observed in Bmal1^-/-^ cells we first annotated mis-expressed elements using a karyotype-like representation of their genomic position. Over-expressed genes and MERVL copies in Bmal1^-/-^ cells were located throughout all chromosomes but tend to cluster at some genomic positions where genes and MERVLs were co-ordinately activated (Figure S3E). For example, transcriptional induction of the six Zscan4 genes occurred concurrently to the activation of three interspersed MERVL copies (Figure S3F). To analyse whether changes in 3D genome organization relates to the concordant transcriptional induction of nearly located MERVLs and genes we compared chromatin organization using chromosome conformation capture ^37^ followed by sequencing (Hi-C) ^38^ in Bmal1^-/-^ and parental JM8^+/+^ cells. Bmal1^-/-^ cells display decreased intra-chromosomal interactions and concomitant increase in inter-chromosomal contacts, demonstrating that BMAL1 may be necessary to maintain an overall chromosome-wide chromatin structure in mESCs (Figure 3D). Next, we asked whether BMAL1 is involved in the compartmentalization of chromosomes in the so-called “A” and “B” compartments (enriched in “active” or “repressive” chromatin respectively). Our analysis indicated that depletion of BMAL1 promotes increased compartment strength (Figure 3E) due to enhanced homotypic interactions between sequences belonging to the A compartment (Figure 3F and 3G). Moreover, a subset of genomic regions (2.14%) transited from the “repressed” B to “active” A compartment upon deletion of Bmal1 (Figure 3G), suggesting that the transcriptional activation of TEs and/or genes induced by BMAL1 depletion was accompanied by the migration of their genomic sequences towards the A compartment. In agreement, 605 MERVL copies (log2FC>1, p<0.05) and 273 protein coding genes (log2FC>1.5, p<0.05) that were transcriptionally induced in Bmal1^-/-^ mESCs showed a tendency to be relocated to the A compartment, as compared to non-induced controls (Figure 3H). In contrast, analysis of the structure of topological associating domains (TADs) in Bmal1^-/-^ and parental JM8^+/+^ cells revealed no major differences (Figure S3G and S3H). We concluded that depletion of BMAL1 in mESCs leads to the transcriptional activation of MERVL retrotransposons and neighbouring genes through a mechanism that involves their repositioning towards chromatin compartment A (Figure 3I), where they may have access to molecular machinery that enables higher rates of RNA transcription.

### Bmal1^-/-^ mESCs display 2C-specific molecular and cellular features

MERVLs and ZSCAN4 are expressed in 2C embryos, and they are downregulated in pluripotent cells ^39, 40^. Thus, augmented expression of these two totipotency markers in Bmal1^-/-^ mESCs cultures (Figure 2E and S3F) suggested that depletion of BMAL1 destabilizes pluripotent cell identity, and cells transit back to a 2CLC state (Figure 4A). In support of this hypothesis, we found that deletion of Bmal1 favours the activation of the 2C transcriptional program (Figure 4B). This includes the upregulation of 2C marker genes such as Zscan4, Dux, Dub1, Tcstv1, Tcstv3 and Eif1a (Figure 4C and S4A), that is translated into a significant increase of ZSCAN4 protein in Bmal1^-/-^ cells compared to JM8^+/+^ (Figure 4D, 4E, 4F) and acquisition of a single enlarged immature nucleolus (Figure 4G and 4H), as previously described for 2C cells ^41^. Moreover, Bmal1^-/-^ cells display enhanced propensity to activate extraembryonic trophectoderm markers (Cdx2, Gata3, Mmp9 and Elf5) than wild-type mESCs (Figure 4I and 4J). Similar results were obtained with independently derived Bmal1^-/-^#2 mESCs (Figure 4C, S4B, S4C, S4D, S4E and S4F). Taken together, we concluded that BMAL1-mediated repression of MERVLs and 2C-associated genes prevents the reversion of pluripotent mESCs towards a 2CLC-state.

**Figure 4.**
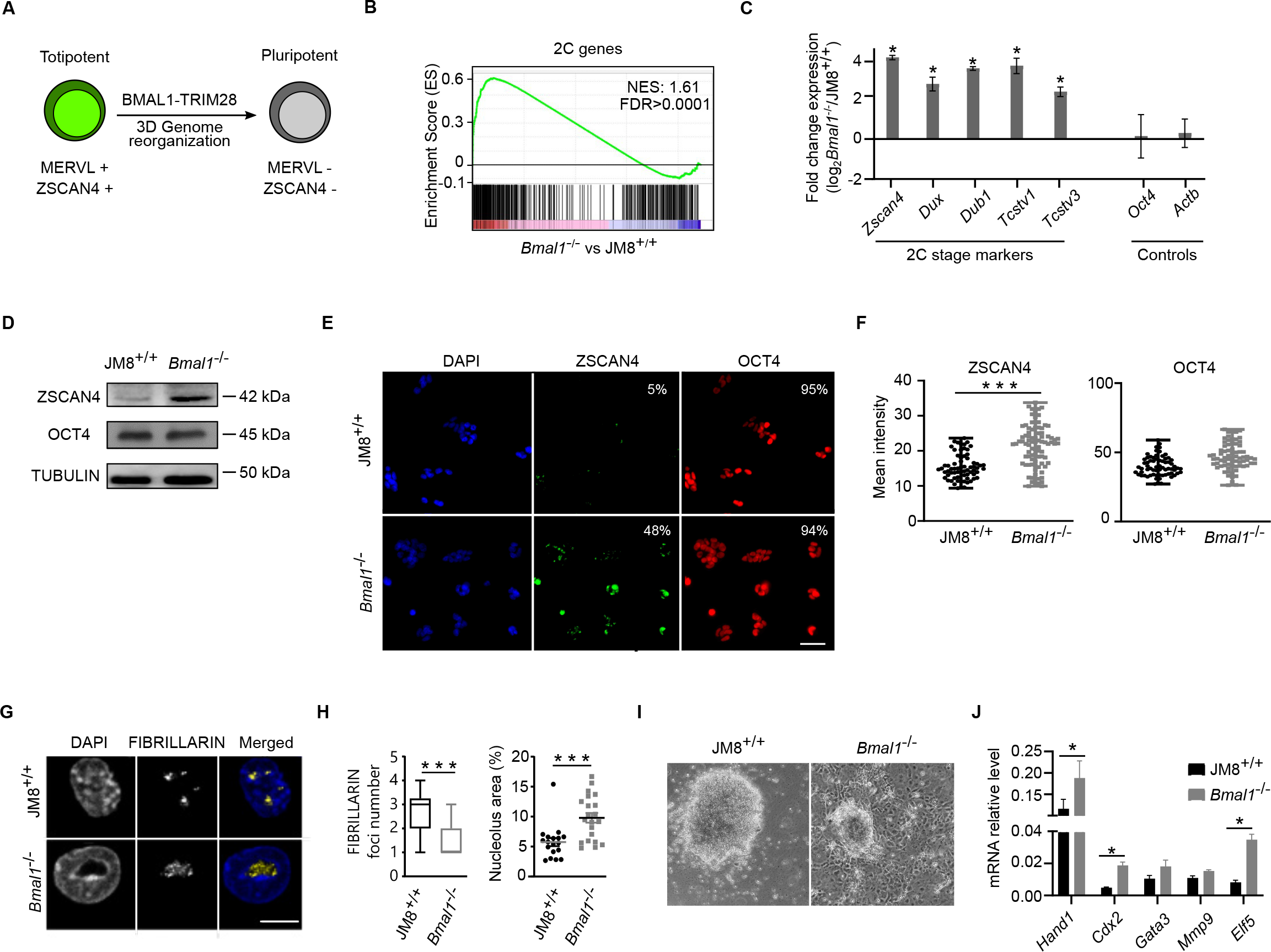
Bmal1^-/-^ mESCs display 2C-specific molecular and cellular features (A) Diagram proposing a mechanism by which BMAL1-TRIM28 facilitates totipotent to pluripotent transition. (B) Gene Set Enrichment Analysis of the 2C-associated gene expression program in Bmal1^-/-^ mESCs. Normalized Enrichment Score (NES) and FDR are indicated. (C) Analysis of mRNA expression by RT-qPCR of 2C-associated genes in Bmal1-/- relative to JM8^+/+^ mESCs. Gene expression was normalized to housekeeping genes (Hmbs, Yhwaz). Oct4 and Actb were included as controls. Mean ± SEM of three experiments is shown. Asterisks indicate p<0.05 in Mann- Whitney test. (D) Western Blot analysis of whole cells lysates showing the expression of ZSCAN4 and OCT4 proteins in JM8^+/+^ and Bmal1^-/-^ mESCs. TUBULIN was used as loading control. (E) Immunofluorescence images of DAPI (blue), ZSCAN4 (green) and OCT4 (red) staining in JM8^+/+^ and Bmal1^-/-^ mESCs. The percentage of cells positive for ZSCAN4 or OCT4 labelling is indicated (n=80?). Scale bar is 100 µm. (F) Plot showing the mean intensity of ZSCAN4 and OCT4 in JM8^+/+^ and Bmal1^-/-^ mESCs (n=80 cells per genotype). Asterisks indicate p<0.0001 in Mann-Whitney test. (G) Immunofluorescence images of the nucleolar protein FIBRILLARIN and DAPI staining in JM8^+/+^ and Bmal1^-/-^ mESCs. Scale bar is 10 µm. (H) Boxplots showing the number of FIBRILLARIN-positive foci per cell (left panel), and the percentage of the area of the nucleus that they occupy (right panel) in JM8^+/+^ and Bmal1^-/-^ mESCs (n=30 cells per genotype). Median values are indicated with horizontal lines. Asterisks indicate p<0.0001 in Mann- Whitney test. (I) Brightfield microscopy images of JM8^+/+^ and Bmal1^-/-^ mESCs after 14 days of growth at low density in ESCs media without ERK or GSK3 inhibitors, but with serum and LIF, that allows heterogenous spontaneous differentiation towards trophectoderm. (J) Gene expression analysis by RT-qPCR of trophectoderm-associated genes in JM8^+/+^ and Bmal1^-/-^ mESCs described in (I). mRNA expression was normalized to Hmbs and Yhwaz housekeeping genes. Mean ± SEM of three experiments is shown. Asterisks indicate p<0.05 in Mann-Whitney test.

### BMAL1-mediated repression of 2CLC-specific features does not require CLOCK protein

We wondered whether BMAL1 requires interaction with CLOCK to repress MERVL elements and 2C- associated genes in mESCs. It has been proposed that CLOCK protein is not expressed in mESCs ^18^, and that the absence of a functional BMAL1-CLOCK heterodimer impedes the production of circadian oscillations in this cell type ^14^. In consonance, we detected very low expression of CLOCK protein in wild type mESCs compared to mouse neural stem cells (NSCs) where the circadian clock is functional (Figure 5A and S5A). We hypothesized that in the absence of the right stoichiometric amount of CLOCK protein, BMAL1 interacts with TRIM28 and carries out a CLOCK-independent alternative function. To challenge this model, we depleted CLOCK protein by generating Clock^-/-^ mESCs using CRISPR/Cas9 (Figure 5B and S5), confirmed pluripotency-associated colony morphology an expression of genes (Figure S5C and S5D), and study whether depletion of CLOCK recapitulated the alterations observed in Bmal1^-/-^ mESCs. In contrast to the phenotype observed in Bmal1^-/-^ cells, the lack of CLOCK protein did not affect the nuclear distribution of TRIM28 protein nor the formation of H3K9me3-labelled heterochromatin foci (Figure 5C). Importantly, mRNA-seq analysis demonstrated that neither MERVL elements, nor any other TE family, was upregulated in Clock^-/-^ cells compared to the JM8^+/+^ cell line (Figure 5D, S5E and S5F). The lack of CLOCK protein resulted in the over-expression of 248 protein coding genes (FC>2, p<0.05), which were mostly unaffected in Bmal1^-/-^ cells (Figure 5E). In addition, 2C-associated genes were not over-expressed in Clock^-/-^ cells (Figure 5F and 5G), the level of ZSCAN4 protein was unaltered (Figure 5H), and the nucleolar organization did not acquire a 2C-typical large nucleolar distribution (Figure 5I and 5J). We concluded that BMAL1 does not require CLOCK protein to repress MERVL TEs nor 2C-associated genes in mESCs. Overall, our results support a model in which interaction between BMAL1 and TRIM28 during prenatal development helps to establish cell-type-specific gene expression programs, while interaction between BMAL1 and CLOCK after birth mediate circadian oscillations of gene transcripts (Figure 5K).

**Figure 5.**
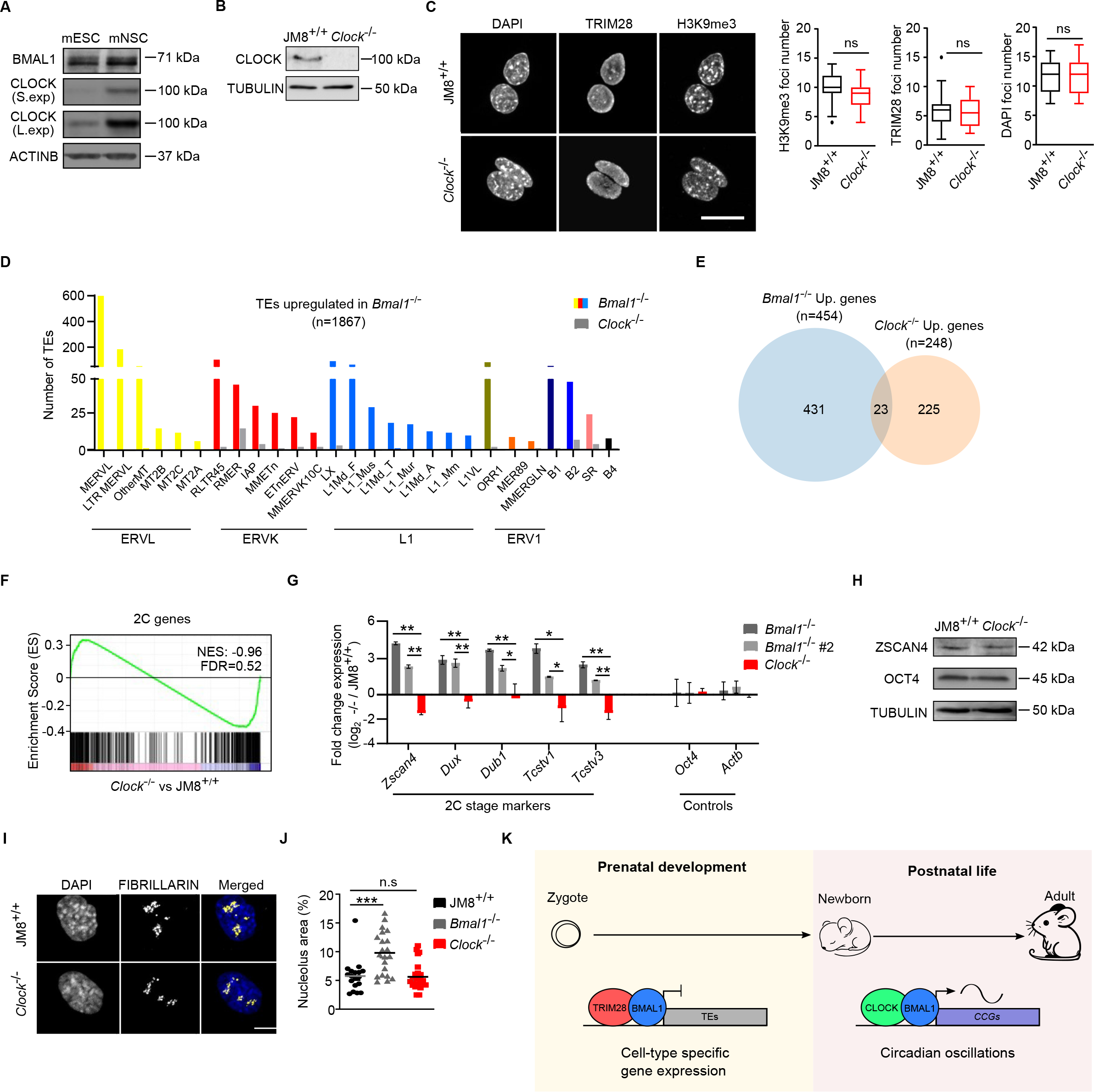
Clock^-/-^ mESCs do not phenocopy Bmal1^-/-^ mESCs (A) Western blot analysis of whole cell extracts comparing the expression of BMAL1 and CLOCK in mESCs and mNSCs. Short (S. exp) and long (L. exp) exposures of the anti-CLOCK signal are presented. ACTINB was used as a loading control. (B) Western blot analysis of whole cell extracts measuring the level of CLOCK protein in JM8^+/+^ and Clock^-/-^ mESCs. TUBULIN is used as a loading control. (C) Immunofluorescence images of DAPI, TRIM28 and H3K9me3 labelling in JM8^+/+^ and Clock^-/-^ mESCs (left panel). Scale bar is 10 µm. Boxplots showing the number of DAPI, TRIM28 and H3K9me3 foci per nucleus (n=50 per genotype) (right panels). (D) Histogram showing the number of TEs overexpressed in Bmal1^-/-^ and Clock^-/-^ mESCs and belong to indicated TE families. (E) Venn diagram showing the overlap between the genes upregulated (FC>2, p-value<0.05) in Bmal1^- /-^ and in Clock^-/-^ mESCs. (F) Gene Set Enrichment Analysis of 2C-associated genes in Clock^-/-^ mESCs. Normalized Enrichment Score (NES) and FDR are indicated. (G) Gene expression analysis by RT-qPCR of 2C-associated genes in Bmal1^-/-^, Bmal1^-/-^#2 and Clock^-/-^ relative to matched parental control mESCs. Expression was internally normalized against housekeeping genes (Hmbs, Yhwaz). Expression of Oct4 and Actb genes were used as controls. Mean ± SEM of three experiments is shown. Asterisks indicate p<0.05 in Mann-Whitney test. (H) Western Blot analysis of whole cells lysates comparing the expression of ZSCAN4 and OCT4 proteins in JM8^+/+^ and Clock^-/-^ mESCs. TUBULIN was used as a loading control. (I) Immunofluorescence images of the nucleolar protein FIBRILLARIN and DAPI staining in JM8^+/+^ and Clock^-/-^ mESCs. Scale bar is 10 µm. (J) Boxplot showing the percentage of the area of the nucleus that is occupied by FIBRILLARIN foci in JM8^+/+^, Bmal1^-/-^, and Clock^-/-^ mESCs (n=30 cells per genotype). Median values are indicated with horizontal lines. Asterisks indicate p<0.0001 in Mann-Whitney test. n.s. indicates non-significant differences. (K) Model proposing a novel non-circadian function of BMAL1 during prenatal development. BMAL1, TRIM28 and CLOCK proteins are depicted bound and regulating transposable elements (TEs) or clock- controlled genes (CCGs).

## Discussion

It is now almost two decades ago that the opposing phenotypes of Clock and Bmal1 mutant mice were reported; altered metabolism and obesity in Clock mutant mice ^8^ versus a drastic progeria-like syndrome in Bmal1^-/-^ adults ^9^. This discrepancy was further exacerbated by a more recent report by the Fitzgerald lab demonstrating that the early aging phenotype of Bmal1^-/-^ mice is mostly due to the function of BMAL1 protein during prenatal development ^10^, when CLOCK is not expressed and circadian oscillations are absent ^14, 18^. Together, these discoveries raised the fundamental question as to what the function of BMAL1 during embryo development is. In this study, we demonstrate that BMAL1 interacts with TRIM28 and facilitates H3K9me3-mediated repression of MERVL TEs in pluripotent cells, and that the loss of BMAL1 function induces a wide-spread transcriptional activation of MERVL elements, 3D genome reorganization and the acquisition of 2C-associated cellular and molecular features. Thus, our findings indicate that during early development BMAL1 is redeployed as a transcriptional repressor of TEs and suggest that this CLOCK-independent function is the key molecular pathway underlying the early aging phenotype observed in Bmal1^-/-^ adult mice.

Our proteomic analysis indicates that BMAL1 interacts with the transcriptional regulators TRIM28 and RIF1 in mESCs, and our functional analyses demonstrate that depletion of BMAL1 impairs TRIM28 binding and H3K9me3 accumulation at MERVL elements, resulting in augmented production of MERVL transcripts. These findings are in consonance with previous reports that support that both TRIM28 and RIF1 can repress transcription of ERVs through SETDB1 and H3K9 methylation ^26, 31, 32, 42^. In addition, the phenotype of Bmal1^-/-^ mESCs is also similar to the one described in RIF1 and SETDB1 null mESCs at the cellular level ^43, 44, 45, 46^, because the three mutant cell lines display overt activation of 2C- associated genes, and higher efficiency of differentiation towards extraembryonic lineages. The main difference between the phenotypes of Bmal1, Trim28 and SetdB1 knockout pluripotent cells is that while the loss of TRIM28 and SETDB1 is incompatible with pluripotency maintenance and early embryogenesis ^32, 47, 48, 49^, the lack of BMAL1 does not fully impair pluripotent self-renewal nor embryo development ^9, 16, 17^. The milder phenotype observed upon Bmal1 deletion is probably a consequence of the recruitment of TRIM28 and SETDB1 to chromatin through BMAL1-independent mechanisms, such as the ones involving ZFPs that contain a Krüppel-associated box domain (KRAB-ZFPs) ^24^. In fitting, while deletion of Trim28 leads to a wide overexpression of TEs of different families in mESCs ^32^, we found that the depletion of BMAL1 mostly influences a specific subset of them.

Although in this study we demonstrate that BMAL1 is required for localization of TRIM28 on chromatin, the details of the molecular interplay between these two proteins are unclear. TRIM28 does not directly bind to DNA, but it is generally recruited to DNA by KRAB-ZFPs ^24^, which bind DNA through their zinc finger domains and recruit TRIM28 through interaction with their KRAB domain ^24^. Because BMAL1 harbours no standalone DNA-binding nor annotated KRAB domains, deciphering the mechanistic details as to how BMAL1 regulates recruitment of TRIM28 to chromatin is complex and will require further studies. In this context it might be relevant to highlight that the BMAL1-CLOCK heterodimer is recruited to E-box-containing promoters through the DNA binding domain that is formed by the two complementary basic helix-loop-helix (bHLH) motifs contained in each of the heterodimer partners ^50^. Thus, it is possible that in the absence of CLOCK protein the standalone bHLH motif of BMAL1 facilitates interactions with MERVL-encoding DNA. Alternatively, BMAL1-TRIM28 might be recruited to MERVL elements through the DNA-binding ZFPs that we detected as BMAL1 interactors in proteomic analyses (ZFP638, ZFP281 and ZFP326).

The expression of MERVLs is strictly regulated during mouse early development, and it is commonly used as a marker of totipotency that is distinctly lost in pluripotent cells ^21^. However, the molecular mechanism by which MERVL elements favour the totipotent state is poorly understood, and it has been suggested that it involves the synthesis of MERVL-chimeric transcripts ^23, 34^, direct modulation of OCT4 and SOX2 levels ^51^, and changes in the 3D organization of the genome ^35, 36^. In this context we found that transcriptional activation of MERVL copies and 2C-associated genes in Bmal1^-/-^ mESCs is not induced by altered levels of OCT4 protein nor production of MERVL-containing chimeric transcripts, but rather through a in cis co-regulation mechanism that involves changes in 3D genome organization. This conclusion is sustained by our Hi-C analysis showing repositioning of transcriptionally induced MERVL elements and protein coding genes towards the A compartment. In addition, in contrast to previous reports analysing 2C embryos ^52, 53, 54^ or CAF-1-depleted 2CLCs ^36^, our analyses revealed no genome-wide decrease of insulation nor formation of new insulation borders due to MERVL activation. Thus, we propose that although full reprogramming of pluripotent chromatin towards totipotency might involve global changes in insulation, formation of MERVL-associated TADs and changes in chromatin compartments, the impact of BMAL1 deletion only induces specific changes in compartmentalization. Consequently, Bmal1^-/-^ mESCs seem to reside in an intermediate state between pluripotency and totipotency, characterized by simultaneous expression of MERVL elements and 2C-specific genes together with a high level of the pluripotency-associated factor OCT4.

Independently of the molecular details explaining the regulatory relationship between MERVL and 2C- asssociated genes, our results provide solid evidence that the transcriptional repression of MERVL elements by BMAL1-TRIM28 is functionally independent of CLOCK. This suggest that the low expression of CLOCK protein observed during mouse development ^18^ facilitates non-circadian alternative interactions between BMAL1 and H3K9me3-related proteins (TRIM28 and RIF1) during embryogenesis. However, we cannot rule out that BMAL1 represses TEs through interaction with TRIM28 in other contexts in which CLOCK protein is present and circadian expression is taking place, because a fraction of BMAL1 monomers might not form activating complexes with CLOCK and would interact with TRIM28 instead. In consonance, a recent study has shown that BMAL1 interacts with TRIM28 to repress the transcription of LINE1 TEs in human mesenchymal stem cells (hMSCs) derived from adult bone marrow ^55^.

To conclude, we propose that BMAL1 interacts with TRIM28 and represses the transcription of TEs during prenatal development, and that this CLOCK-independent function constitutes the molecular basis of the early aging phenotype of Bmal1^-/-^ adult mice ^9, 10^. This idea is in agreement with emerging reports that indicate that the activity of retrotransposons is an important source of aging in metazoan organisms through the alteration of gene expression programs and DNA integrity ^56, 57^. Thus, future research will have the interesting challenge of dissecting the molecular details as to how unintended expression of TEs during prenatal life might lead to premature aging during adulthood. Additionally, it is possible that the progeria-like syndrome of Bmal1^-/-^ mice is based on a failure to maintain genome stability during development due to TE-independent mechanisms, because the BMAL1-interacting factors TRIM28 and RIF1, and H3K9me3, are required for effective DNA repair and/or correct chromosome segregation ^58, 59, 60^.

## Methods

### Cell culture and derivation of mESCs lines

JM8^+/+^ (background C57BL6/N)-derived mESCs were grown on 0.1% gelatin-coated dishes at 37°C and 5% CO2, and in N2B27 serum-free media containing Mitogen-Activated Protein Kinase (MEK) inhibitor PD0325901 (1 μM) (Millipore), Glycogen Synthase Kinase 3 (GSK3) inhibitor CHIR99021 (3 μM) (Millipore) and Leukaemia-Inhibitory Factor (LIF) as previously described ^61^. Neural Stem Cells (NSCs) were derived from wild type JM8 mESCs using a described protocol ^62^.

JM8-derived Bmal1^-/-^ mESCs were previously described ^16^. Clock-/- mESCs were derived from JM8^+/+^ mESCs following a described protocol ^16^. Briefly, the Clock gene was targeted by CRISPR/Cas9 on exon 6 using guide RNAs cloned in PX458, lipofectamine transfection followed by flow cytometry sorting and isolation of genetic clones. Genetic edition of targeted exon was analysed by PCR (oligos Table S3) and sanger sequencing followed by the ICE CRISPR analysis tool (Synthego). The absence of CLOCK protein was checked by western blot in clones harbouring nonsense mutations.

To derive JM8 cells that express two FLAG sequences in tandem (2XFLAG) on the N-terminus of BMAL1 protein we obtained Bmal1 cDNA from JM8 cells, fused it with a 2XFLAG sequence, cloned it into the pCAG-Puromycin expressing vector, and transfected into cells to obtain genetically clonal populations of mESCs. Specifically, we isolated mRNA from JM8^+/+^ cells using Trizol reagent (Thermofisher), reverse transcribe it using SuperScript III Reverse Transcriptase (Invitrogen) and specific primers against Bmal1 mRNA. Forward primer (Table S3) anneals on the second codon of Bmal1 mRNA and include an adapter stuffer sequence on 5’ end. Reverse primer (Table S3) anneals on the stop codon of Bmal1 and include a NotI target sequence on the 3’ end. A 2XFLAG sequence containing a NotI restriction site on the 5’ and the stuffer sequence on the 3’ was amplified with KAPA Hifi HotStart Readymix (Kapa Byosystems) PCR system, using the pCAG-2XFLAG-JARID2-Puromycin plasmid as template ^63^, a forward primer with NotI adapter sequence (Table S3) and a reverse primer annealing on the stuffer sequence (Table S3). 2XFlag:Bmal1 DNA molecules were generated by overlap extension PCR using the KAPA Hifi HotStart Readymix, the forward primer designed to amplify 2XFLAG sequence and the reverse primer designed to amplify the Bmal1 cDNA. PCR fragment was digested using NotI (NEB) and ligated with NotI-digested pCAG-Puromycin plasmid using T4 DNA ligase (NEB) to generate the pCAG-2XFLAG- BMAL1 plasmid. Sequence fidelity of the 2XFlag:Bmal1 transgene was confirmed by PCR followed by Sanger sequencing with multiple oligos covering all the open reading frame (Table S3). JM8^+/+^ mESCs were transfected with pCAG-2XFLAG-BMAL1 plasmid using lipofectamine 2000 and cells resistant to 0.3 µg/mL Puromycin were selected for 10 days. Resistant colonies were isolated and the expression of FLAG-BMAL1 was measured by western blot to select clones in which the level of expression of FLAG-BMAL1 and endogenous BMAL1 were similar.

### Analysis of spontaneous differentiation towards trophectoderm

To measure spontaneous differentiation of JM8^+/+^ and Bmal1^-/-^ mESCs towards trophectoderm, cells were transferred to serum mESC media without MEK and GSK3 inhibitors as described previously ^46^. Briefly, cells were plated at low density (100 cells/cm^2^) and allowed to form colonies during ten days on 0.1% gelatin-coated dishes with DMEM KO (Gibco) media supplemented with 10% FBS (Gibco), LIF, penicillin/streptomycin (Gibco), L-glutamine (Gibco), 2-mercaptoethanol (Gibco).

RT-qPCR analysis

Total RNA was extracted using Trizol reagent (Thermofisher), digested with DNAseI (Invitrogene) and reverse transcribed using RevertAid Frist Strand cDNA synthesis kit (Thermofisher). For the analysis of expression of MERVL elements the RNA was additionally digested with RQ1 DNAse (Promega) to guarantee the absence of contaminant genomic DNA. qPCR was carried out using GoTaq qPCR Master Mix with SYBR Green (Promega). Primers used are provided in table S3. Primers used for expression of pre-rRNA transcripts were described previously ^64^.

### Immunofluorescence analysis

Immunofluorescence analysis was carried out as described previously ^65^. Briefly, mESCs were fixed for 20 minutes in 2% paraformaldehyde, permeabilized 5 minutes in 0.4% Triton-X100 and blocked for 30 minutes in blocking buffer (PBS supplemented with 0.05% Tween 20, 2.5% bovine serum albumin and 10% goat serum). Detailed information of primary antibodies used is included in Table S3. Goat anti-mouse Alexa fluor 488 (Thermofisher, A-11001), goat anti-rabbit Alexa fluor 555 (Thermofisher, A- 21429) were used as secondary antibodies. Vecta-shield mounting media with 1 µg/ml freshly added DAPI was used. Slides were analysed using a widefield fluorescence microscope Zeiss Axio Imager and the Image J software.

### Western blot analysis

Western blots of whole cell extracts or histone preparations were carried out as previously described^16^. Primary antibodies used are listed in Table S3. A secondary specie-specific antibodies conjugated to horseradish peroxidase was used (anti-rabbit-HRP, GE-Healthcare). Clarity Western ECL reagents (Bio-Rad) was used as chemiluminescent substrate.

### Co-immunoprecipitation followed by western blot and mass spectrometry analysis

Nuclear extracts of around 4x10^7^ FLAG-BMAL1 and JM8^+/+^ cells were prepared as previously described ^66^. Nuclear extracts were quantified with Bradford and 2mg of lysate was mixed with 6 µg of antibody and 50 µL of Protein G magnetic beads (Dynabeads, Invitrogen) in 500 µl of buffer C (300 mM NaCl, 20 mM Hepes-KOH, pH 7.9, 20% v/v glycerol, 2 mM MgCl2, 0.2 mM EDTA, 0.1% NP-40, protease inhibitors, and 0.5 mM dithiothreitol) overnight. The reaction was washed 4 times: twice with buffer C and 0.5% NP-40, followed by one with PBS and 0.5% NP-40 and a final one with PBS. To perform western-blot analyses, immunoprecipitated proteins were eluted in 70 μl of NuPage loading buffer (final concentration of loading buffer 2X and 50 mM DTT) per IP at 70°C for 15 min.

For mass spectrometry analysis the magnetic beads used in immunoprecipitation were cleaned three times with 500 µl of 200 mM ammonium bicarbonate and 60 µl of 6M Urea / 200mM ammonium bicarbonate were added. Samples were then reduced with dithiothreitol (30 nmol, 37 °C, 60 min), alkylated in the dark with iodoacetamide (60 nmol, 25 °C, 30 min) and diluted to 1M urea with 200 mM ammonium bicarbonate for trypsin digestion (1 µg, 37°C, 8h, Promega). After digestion, peptide mix was acidified with formic acid and desalted with a MicroSpin C18 column (The Nest Group, Inc) prior to LC-MS/MS analysis. The samples were then analysed using a LTQ-Orbitrap Fusion Lumos mass spectrometer (Thermo Fisher Scientific, San Jose, CA, USA) coupled to an EASY-nLC 1200 (Thermo Fisher Scientific (Proxeon, Odense, Denmark). Peptides were loaded directly onto the analytical column and were separated by reversed-phase chromatography using a 50-cm column with an inner diameter of 75 μm, packed with 2 μm C18 particles. Chromatographic gradients started at 95% buffer A and 5% buffer B with a flow rate of 300 nl/min and gradually increased to 25% buffer B and 75% A in 52 min and then to 40% buffer B and 60% A in 8 min. After each analysis, the column was washed for 10 min with 100% buffer B. Buffer A: 0.1% formic acid in water. Buffer B: 0.1% formic acid in 80% acetonitrile. The mass spectrometer was operated in positive ionization mode with nanospray voltage set at 2.4 kV and source temperature at 305°C. The acquisition was performed in data-dependent acquisition (DDA) mode and full MS scans with 1 micro scans at resolution of 120,000 were used over a mass range of m/z 350-1400 with detection in the Orbitrap mass analyser. Auto gain control (AGC) was set to ‘standard’ and injection time to ‘auto’. In each cycle of data-dependent acquisition analysis, following each survey scan, the most intense ions above a threshold ion count of 10000 were selected for fragmentation. The number of selected precursor ions for fragmentation was determined by the “Top Speed” acquisition algorithm and a dynamic exclusion of 60 seconds. Fragment ion spectra were produced via high-energy collision dissociation (HCD) at normalized collision energy of 28% and they were acquired in the ion trap mass analyzer. AGC and injection time were set to ‘Standard’ and ‘Dynamic’, respectively and isolation window of 1.4 m/z was used. Digested bovine serum albumin (New England Biolabs) was analysed between each sample to avoid sample carryover and to assure stability of the instrument. QCloud has been used to control instrument longitudinal performance during the project.

Acquired spectra were analysed using the Proteome Discoverer software suite (v1.4, Thermo Fisher Scientific) and the Mascot search engine (v2.6, Matrix Science). Data was analysed against a Swiss- Prot mouse database (as in March 2021, 17082 entries) plus a list of common contaminants and all the corresponding decoy entries. For peptide identification a precursor ion mass tolerance of 7 ppm was used for MS1 level, trypsin was chosen as enzyme and up to three missed cleavages were allowed. The fragment ion mass tolerance was set to 0.5 Da for MS2 spectra. Oxidation of methionine and N-terminal protein acetylation were used as variable modifications whereas carbamidomethylation on cysteines was set as a fixed modification. False discovery rate (FDR) in peptide identification was set to a maximum of 5%. SAINT express algorithm was used to score protein-protein interactions and STRING software was used to identify functional clusters among FLAG-BMAL1-interacting proteins.

### Calibrated chromatin immunoprecipitation sequencing (cChIP-seq) and data analysis

cChIP-seq were carried out as previously described ^67^ with some modifications. Briefly, cells were trypsinized, resuspended in mESCs cell-culture media and counted. mESCs cells were mixed with human A549 cells in a proportion of 25:1 as spike-in control per reaction. Four million JM8^+/+^ or Bmal1^-/-^ mESCs were used for each cChIP-seq reaction. Ten million cells were used as starting material in the FLAG-BMAL1 ChIP-seq that did not require calibration. Cells were spined and resuspended in 37°C complete media at a density of 5x10^6^ cells/ml and incubated in a rotating platform for 12 min with 1% formaldehyde at room temperature. After quenching formaldehyde fixation with glycine (125mM final concentration), cells were resuspended in swelling buffer (25 mM HEPES pH 7.9, 1.5 mM MgCl2, 10 mM KCl, 0.1% NP-40) at a density of 2.5x10^6^ cells/ml. Nuclei were isolated using a Dounce homogenizer (tight pestle; 50 strokes). Nuclei were resuspended in sonication buffer (1x10^7^ cells/ml) and sonicated 1:30h full power 4°C (30sec ON /30sec Off). Primary antibodies (Table S3) were added to chromatin and incubated in a rotating wheel at 4°C overnight. Protein G magnetic beads (Dynabeads, Invitrogen) were added, and chromatin was incubated for 5 hours. Washes were carried out for 5 min at 4°C with 1 ml of the following buffers: 1× sonication buffer, 1× wash buffer A (50 mM Hepes (pH 7.9), 500 mM NaCl, 1 mM EDTA, 1% Triton X-100, 0.1% Na-deoxycholate, and 0.1% SDS), 1× wash buffer B (20 mM tris (pH 8.0), 1 mM EDTA, 250 mM LiCl, 0.5% NP-40, and 0.5% Na-deoxycholate), and 2× TE buffer (pH 8). DNA was eluted in elution buffer (50 mM Tris pH 7.5, 1 mM EDTA, 1% SDS) and reversed cross-linked overnight at 65°C in 160mM NaCl and 20 µg/ml RNase, followed by 2h 45°C incubation with proteinase K (220 µg/ml).

Libraries of immunoprecipitated DNA were generated from 3 ng of starting DNA with the NEBNext Ultra DNA Library Prep kit for Illumina (New England Biolabs) according to manufacturer’s instructions at the CRG Genomics Core Facility (Barcelona) and sequenced using a NextSeq 500 Illumina technology. 20–30 million reads (75 bp paired-end-reads) were obtained for each library.

Quality of libraries were evaluated using FastQC v0.11.5 software. Reads were aligned using Bowtie2 ^68^ to the genome sequence of the concatenated mouse (mm10) and spike-in genomes (hg19) using the “–no-mixed” and “–no-discordant” options. Unique-mapping and randomly allocated multi-hit reads were selected using bowtie2 (Random mode, without post filtering the results, and keeping alignments when alignment score was higher than the second valid alignment)^69^. Duplicates were then removed and the alignments to the mouse genome were separated from the human spike-in. The mouse reads were sorted and indexed using SAMTools and sambamba ^70, 71^. Mouse reads were then randomly subsampled using the calculated down sampling factor for each ChIP using a random seed of 123. Down sampling factor was calculated for each sample considering the ratio of sequences aligned to the inputs of the spike-in and the mouse genome, as well as the number of reads aligning to the spike-in of each ChIP sample. BigWigs were generated using the deepTools suite ^72^ without further normalization for calibrated samples and with reads per kilobase per million (RPKM) for uncalibrated ones. Peak calling was performed with MACS3 ^73^ using FDR< 0.05 as threshold. The genome location of detected peaks was classified using HOMER ^74^.

Read coverage of the cChIP-seqs was calculated with the R-package CoverageView version 1.38.0. with no further normalization. FLAG-BMAL1 non-calibrated ChIP-seq coverage plots was measured as reads per million (RPM). TRIM28 peaks with average read signal above the 95th percentile or below the 5th percentile were discarded, and average read signal around the centre of the remaining peaks was plotted. Binning of 10 bp was used. Read coverage of upregulated MERVLs or solo LTR sequences was calculated by dividing the DNA sequence of each element in 100 windows, calculating the average reads of each window, and plotting the average value of all elements per window. Average read coverage around the TSS of annotated genes was calculated. To analyse the read coverage in contiguous genomic intervals, the genome sequence was divided in contiguous windows of 1000 bps and total reads mapping to these windows was calculated. The sum of ten consecutive 1000 bp windows was calculated for TRIM28, BMAL1 and FLAG-BMAL1 ChIP-seqs, while the sum of 100 consecutive 1000 bp windows was used to analyse H3K9me3 distribution. These values were used to generate correlation plots and histograms with number of windows. Coverage values for H3K9me3 cChIP-seq was normalized by subtracting input signals.

Reads signal along the consensus sequences of MERVLs and LTRs was obtained by aligning reads to the consensus sequences retrieved from RepBase (www.girinst.org/repbase/) using Samtools ^70^. The average coverage of consensus sequences from different families of TEs was calculated. The fold change between JM8^+/+^ and Bmal1^-/-^ was calculated and plotted as red-white heatmap.

### mRNA sequencing and data analysis

Biological replicates of total RNA from 2x10^5^ JM8^+/+^ and Clock^-/-^ mESCs growing in 2i + LIF media using Trizol reagent (Thermofisher) were obtained. mRNA library preparation using TruSeq Stranded mRNA kit (Illumina) and Illumina sequencing (30 million reads, 75-bp paired end) was carried out at National Centre for Genomic Analysis - Centre for Genomic Regulation (CNAG-CRG). Datasets of published mRNA-seq comparing JM8^+/+^ and Bmal1^-/-^ mESCs ^16^ were included in downstream analyses.

Quality of mRNAseq libraries was evaluated using FastQC v0.11.5 software ^75^. RNA paired-end reads were aligned to mm10 mouse genome assembly with STAR v2.5.3a ^76^ and quantified with featureCounts v2.0.1 ^77^ using Gencode vM22 annotation. Differential expressed genes were determined with the R package DESeq2 v1.36 ^78^. Shrinkage of effect size was performed on DESeq2 results using the apeglm method through the function lfcShrink. Gene Set Enrichment Analysis (GSEA) was performed with the GSEA software from the Broad Institute and UC San Diego ^79^. The set of genes expressed in embryos during the 2 cell stage and used in GSEA analyses was obtained from previous studies ^80^.

### Analysis of expression of transposable elements

For Transposable Elements (TE) expression analysis the SQuIRE v0.9.9.92 pipeline ^81^ was used to count transposable elements expression following default parameters. Code was modified to include simple repeats, satellites, and low complexity regions in this analysis. TE differential expression analysis was performed from SQuIRE counts using DESeq2 as previously stated. mm10 repeat annotation (RepeatMasker) was downloaded from the UCSC table browser.

Analysis of the expression of different of MERVL elements in Bmal1^-/-^ mESCs was performed by calculating the fold change expression relative to JM8^+/+^ cells in group of elements described elsewhere ^82^. Briefly, the MERVL-int (>5kb) elements flanked by two MT2_Mm or MT2C_Mm were categorized as full-length elements. Additionally, MT2 and MERVL-int were required to be in the same strand, and the total length (2xMT2+int) should be < 10kb. MT2 copies lacking nearby MERVL-int (>5kb) were classified as solo MT2. The remaining copies were defined as other MERVL copies.

Identification of the nearest protein coding genes annotated around the TEs that are differentially expressed in Bmal1^-/-^ mESCs was achieved by using annotatePeakInBatch function in ChIPpeakAnno R package ^83^.

Quantification of chimeric transcripts was performed using ChimeraTE ^84^. Chimeric transcripts formed by elements annotated in the reference genome were detected using ChimeraTE mode 1 considering only genes with TE copies located 5 kb upstream or downstream. A genome-blind approach was additionally performed using ChimeraTE mode 2 to detect chimeras from fixed and polymorphic TEs without the reference genome. Reference genome, reference transcripts and gene annotations files were obtained from GENCODE assembly GRCm38.p6 (mm10). TE annotations and TE reference transcripts in the mouse genome (RepeatMasker 4.0) was downloaded from UCSC web page.

MERVL elements overexpressed in Bmal1^-/-^ cells relative to parental cells were classified as self-dependent (intergenic or within a gene that is not overexpressed in Bmal1^-/-^ cells) or gene-dependent (within a gene that is overexpressed (FC>2, p<0.05) in Bmal1^-/-^ cells) was carried out described previously ^85^ with some modifications. To annotate the position of induced TEs relative to genes (mm10 annotations from NCBI RefSeq database downloaded from UCSC Table Browser ^86^), the R package ChIPpeakAnno ^83^ was used with the following genomic priority: 5’ UTR > 3’ UTR > Exon > Intron > Intergenic. In addition, TEs that fall within a gene that is not expressed in neither JM8^+/+^ nor Bmal1^-/-^ cells were considered as self-dependent.

Karyotype-like analysis of the genome-wide distribution of deregulated genes and MERVL elements was performed using karyoploteR ^87^.

### Analysis of mRNA alternative splicing variants

Analysis of mRNA alternative splicing variants was performed in Bmal1^-/-^ (relative to parental JM8^+/+^ cells). A previously described positive control was used (shZfp207 versus parental control mESCs ^88^). Mouse genomic sequences (mm10) and gene annotations (refSeq) were retrieved from the UCSC genome browser. Mappings to the mouse genome were carried out with STAR v2.7.3 ^76^ using the SQUIRE ^81^ optimized parameters (--outFilterMultimapNmax 500 --winAnchorMultimapNmax 500 -- alignEndsProtrude 100 DiscordantPair --outFilterScoreMinOverLread 0.4 -- outFilterMatchNminOverLread 0.4 --chimSegmentMin 17 --alignIntronMax 300000 -- outSAMattrIHstart 0). rMAts v4.1.1 89 was used for alternative splicing analyses with options (--allowclipping --variable-read-length --readLength 100 -t paired) and results were gathered (FDR<0.05) for 5 classes of events: SE, MXE, A5SS, A3SS and RI.

### Chromosome conformation capture analyses by Hi-C

Hi-C was performed as previously described ^90^ with minor modifications. In brief, 4x10^6^ JM8^+/+^ and Bmal1^-/-^ cells were fixed with formaldehyde 1% and digested overnight at 37°C using 400U of MboI (New England Biolabs, #R0147M). After filling with 50nM biotin-dATP (Invitrogen, #10484552) and 50U of Klenow polymerase (New England Biolabs, #M0210M), proximity ligation was carried out overnight at 16°C with 10000U of T4 DNA Ligase (New England Biolabs, #M0202M). Chromatin was reverse-crosslinked with 16U of Proteinase K (New England Biolabs, #P8107S) and 100 ug RNAase A (Thermo Scientific, #EN0531), DNA was purified using AMPure XP beads (Beckman Coulter, #A63881) and sonicated using Sonicator Bioruptor PICO (Diagenode) to produce fragments of 300-400 bp. Sonicated fragments containing biotin were immobilized on MyOne Streptavidin T1 beads (Invitrogen, #65601), end-repaired with the NEBNext End Repair Module (New England Biolabs, #E6050L) and A-tailed with the NEBNext dA-Tailing Module (New England Biolabs, #E6053L). After adapters ligation, libraries were indexed and amplified with NEBNext UDI primer pairs (New England Biolabs, #E6440S) during 8 PCR cycles with NEBNext HiFi PCR Master mix (New England Biolabs, #M0541S). Fragments of DNA 300 and 800 bp were then using AMPure XP beads (Beckman Coulter, #A63881). Sequencing was performed at National Centre for Genomic Analysis (CNAG) using NovaSeq 6000 technology and sequencing >400 millions of 150 bp PE reads per sample.

Hi-C datasets were processed using TADbit ^91^. Specifically, for both JM8^+/+^ and Bmal1^-/-^ cells, paired-end FASTQ files of 2 Hi-C replicates, previously assessed for reproducibility ^92^, were merged and mapped to mouse GRCm38/mm10 reference genome applying a fragment-based iterative strategy ^93^ using the GEM mapper ^94^. Mapped reads were filtered using TADbit with default parameters, which removed self-circles, dangling ends, duplicated and random breaks among other minor artefactual reads ^91^. After mapping and filtering, the resulting Hi-C matrices contained a total of 505,149,739 valid pairs for JM8^+/+^ and 285,113,464 for Bmal1^-/-^ mutant cells. The resulting raw Hi-C interaction matrices were next normalized with ICE balancing ^93^ at the resolutions of 10kb, 50Kb, 100Kb, 500Kb, and 1Mb.

Compartment analysis was performed on observed-over-expected contact maps at resolutions equivalent to 20kb, 50kb, and 100kb bins using the cooltools eigs-cis module ^95^. Active (A) and inactive (B) compartment types were assigned by GC-content. Saddle plots were generated using the cooltools saddle module. Compartment strength was calculated as the ratio of homotypic (AA+BB) over heterotypic (AB+BA) compartment contacts. The top 20 % of observed/expected values for both homotypic and heterotypic interactions were chosen. Insulation scores were calculated using cooltools insulation module at resolution of 50kb, which identified boundaries categorized into strong and weak as implemented in the module.

## General

R version 4.2.2 and R-studio version 2023.6.1.524 was used. Results were plotted with the ggplot2 package version 3.4.2. GraphPad Prism9 was used for statistical analysis and data presentation.

## Data access

Datasets are available at GEO-NCBI with accession number GSE263285. Raw proteomics data has been deposited in PRIDE repository with identifier PXD050769. Datasets will be made public upon publication acceptance.

## Acknowledgements

We thank core facilities at the Center for Genomics and Oncological Research (GENYO) including the cytometry and microscopy units for excellent technical assistance. We also thank the Centro Nacional de Análisis Genómico (CNAG) and the proteomics facility at the Center for Genomic Regulation (CRG) for support with ChIP-seq, mRNA-seq and proteomic experiments. The Landeira lab is supported by the Spanish ministry of science and innovation (PID2019-108108-100, EUR2021-122005, PID2022- 137060NB-I00), the Instituto de Salud Carlos III (IHRC22/00007), the Andalusian regional government (PIER-0211-2019, PY20_00681) and the University of Granada (A-BIO-6-UGR20) grants. The Marti-Renom lab acknowledges support from the Spanish ministry of science and innovation (PID2020- 115696RB-I00) as well as the Catalan Government through the AGAUR agency (SGR 01127). The work in Sara R. Heras lab was supported by the Spanish ministry of science and innovation (PID2020- 115033RB-I00, RYC-2016-21395, CNS2023-145402) and the Andalusian regional government (PY20_00619 y A-CTS-28_UGR20) grants. We thank the IMPULSE visitor program Severo Ochoa 2022 for funding Amador Gallardo to carry out chromosome conformation capture experiments at the CRG. Efres Belmonte Reche was supported by a Maria Zambrano fellowship financed by NextGenerationEU/European Union.

## Author contributions

DL designed and conceptualized the study. AG, MM-M, JD-R, LL-O and PT-R designed, performed, and analysed bench experiments. EB-R, GP, IF-R, NB and MAM-R performed bioinformatic analyses. SR-H, AS-P, AM-E, and MAM-R provided scientific advice and resources. AG and DL wrote the manuscript. All authors provided scientific feedback. DL obtained funding and supervised research.

---

**Figure S1.**
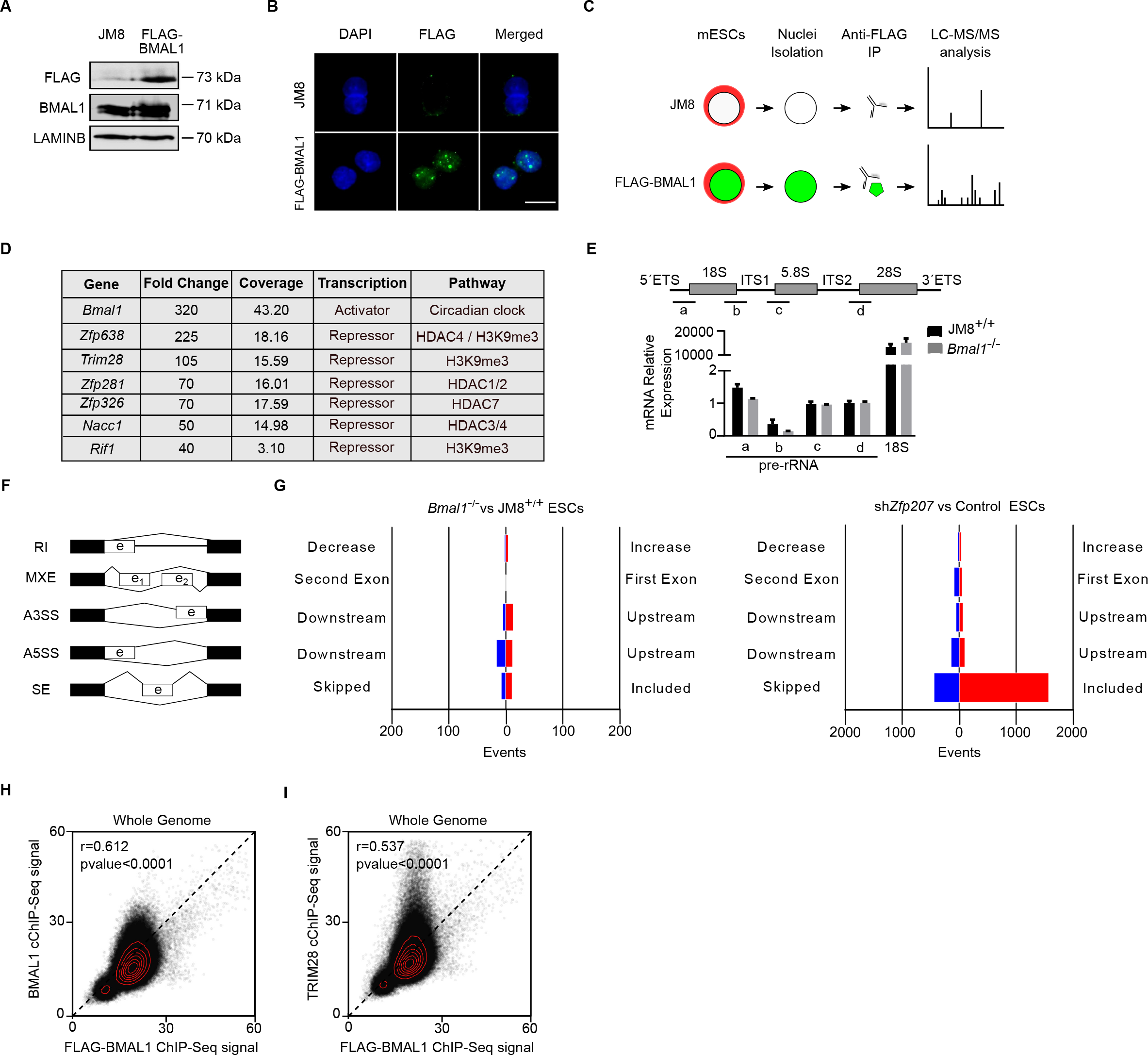
BMAL1^-/-^ mESCs display unaltered expression of mRNA splicing variants and rRNA production (A) Western blot analysis of whole cell lysates using indicated antibodies in JM8 and FLAG-BMAL1 expressing mESCs. Anti-LAMINB was used as loading control. (B) Immunofluorescence analysis of JM8 and FLAG-BMAL1 mESCs using an anti-FLAG antibody (green) and DAPI staining (blue). Scale bar represents 10um. (C) Schematic diagram of the approach used to identify biochemical interactors of BMAL1 in mESCs. Nuclear extracts of FLAG-BMAL1-expressing and parental JM8 control cells were subjected to anti-FLAG immunoprecipitation followed by liquid chromatography and tandem mass-spectrometry. (D) Table summarizing transcriptional regulators interacting with FLAG-BMAL1, as identified by the STRING algorithm. Fold change abundance in FLAG-BMAL1 relative to control JM8 cells, coverage of the protein and information about their annotated role in transcriptional regulation are shown. (E) RT-qPCR analysis of the expression of pre-rRNA transcript in JM8^+/+^ and Bmal1^-/-^ mESCs. The position primers used to measure pre-rRNA transcript is indicated in the upper diagram (a, b, c, d). (F) Schematic representation of the alternative splicing events analysed by mRNA-seq. (G) Bar plots comparing the distribution of alternative splicing changes in Bmal1^-/-^ (left panel) and shZfp207 positive control (right panel) mESCs relative to matched parental cells. (H) Plot showing the genome wide correlation between BMAL1 and FLAG-BMAL1 binding in ChIP-seq experiments in mESCs. The genome was divided in adjacent 10kb-sized windows for which total reads were calculated and plotted. Pearsońs correlation coefficient (r) and p-value are indicated. (I) As in (H) but the for the signals of TRIM28 and FLAG-BMAL1 ChIP-seq experiments.

**Figure S2.**
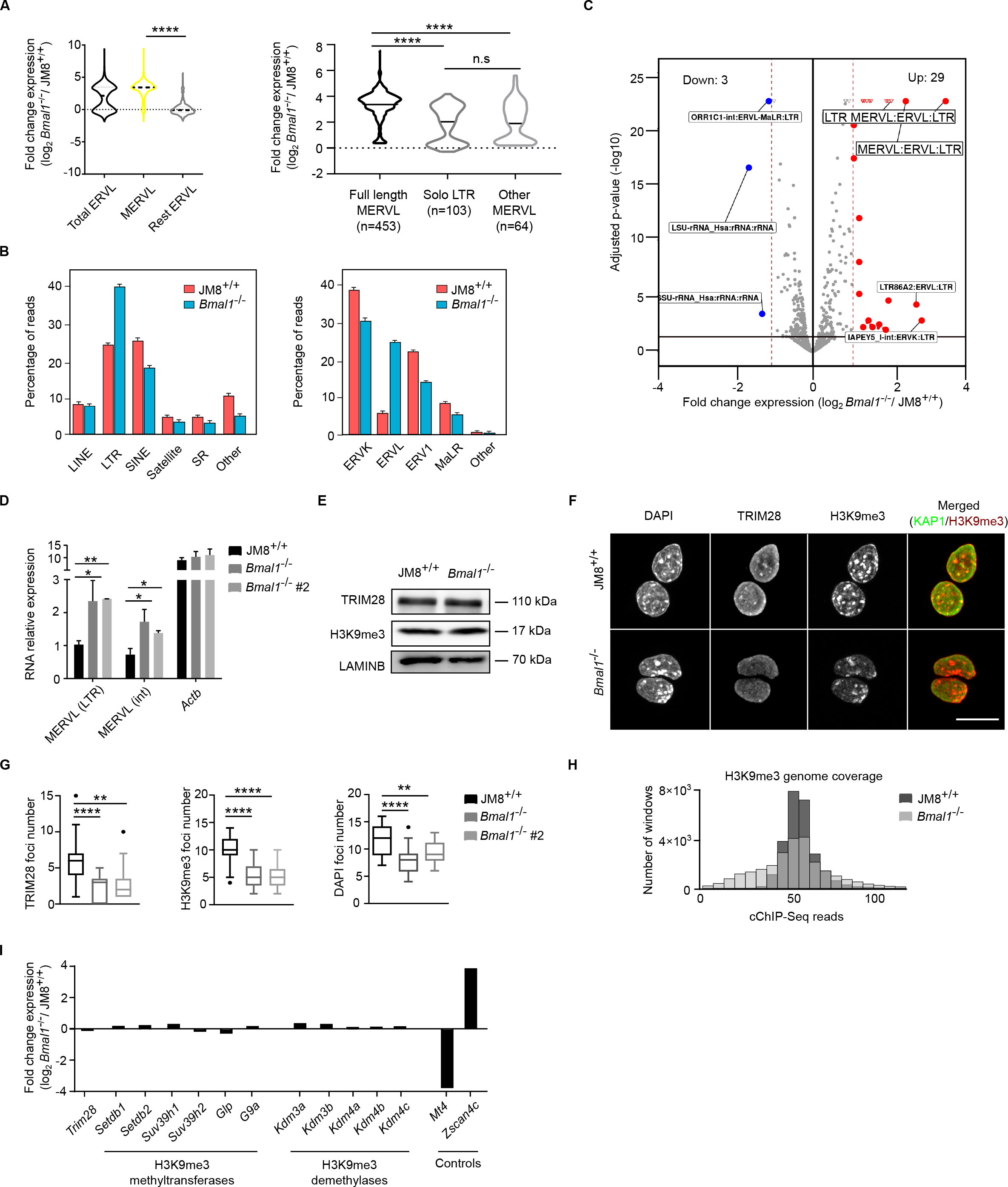
Bmal1^-/-^ mESCs display a partial loss of H3K9me3 on chromatin (A) Violin plots analysing the mRNA fold change expression between Bmal1^-/-^ and JM8^+/+^ cells of all ERVL elements expressed in Bmal1^-/-^ or JM8^+/+^ mESCs (n=2042) (left panel) or full length MERVLs (n=453) (black), solo LTR (n=103) (dark grey) or other MERVLs (n=64) (light grey) (right panel). Discontinuous horizontal lines within each category indicate the median value. Four asterisks indicate p<0.0001 using a Mann-Whitney test. (B) Histogram showing the expression of TEs elements in JM8^+/+^ and Bmal1^-/-^ mESCs assayed by mRNAseq. Percentage of reads mapping to each category are shown. (C) Volcano plot showing the families of TEs that are upregulated (FC>2, p-value<0.05, red dots), downregulated (FC<2, p-value<0.05) or unchanged (grey dots) in Bmal1^-/-^ compared to JM8^+/+^ mESCs. (D) Analysis of the expression of RNA produced by LTR or the internal (int) region of MERVL elements by RT-qPCR analysis in JM8^+/+^, Bmal1^-/-^ and Bmal1^-/-^#2 mESCs. RNA expression is normalized relative to expression of Hmbs and Yhwaz genes. Actb was included as a control. Mean ± SEM of three experiments is shown. Asterisks indicate *p<0.05, **p<0.01 in Mann-Whitney test. (E) Western blot analysis of whole cell extracts comparing global levels of TRIM28 and H3K9me3 in JM8^+/+^ and Bmal1^-/-^ mESCs. LAMINB was used as a loading control. (F) Representative pictures of immunofluorescence experiments using anti-TRIM28, anti-H3K9me3 and DAPI staining in JM8^+/+^ and Bmal1^-/-^ mESCs. Scale bar is 10 µm. (G) Box plots presenting the number of foci positive for TRIM28, H3K9me3 or DAPI signal in JM8^+/+^ and two Bmal1^-/-^ cell lines. Number of foci in fifty nuclei were quantified for each genotype. Asterisks indicate **p<0.01, ****p<0.0001 in Mann-Whitney test. (H) Histogram comparing the genome wide level of H3K9me3 in JM8^+/+^ and Bmal1^-/-^ mESCs. Number of windows for a given total read signal upon dividing the genome in 100kb-sized adjacent windows is plotted. (I) Histogram showing the relative expression of genes involved in the synthesis of H3K9me3 in Bmal1^- /-^ and JM8^+/+^ mESCs. mRNA-seq read counts are plotted. Expression of Mt4 and Zscan4c genes provide differentially expressed controls.

**Figure S3.**
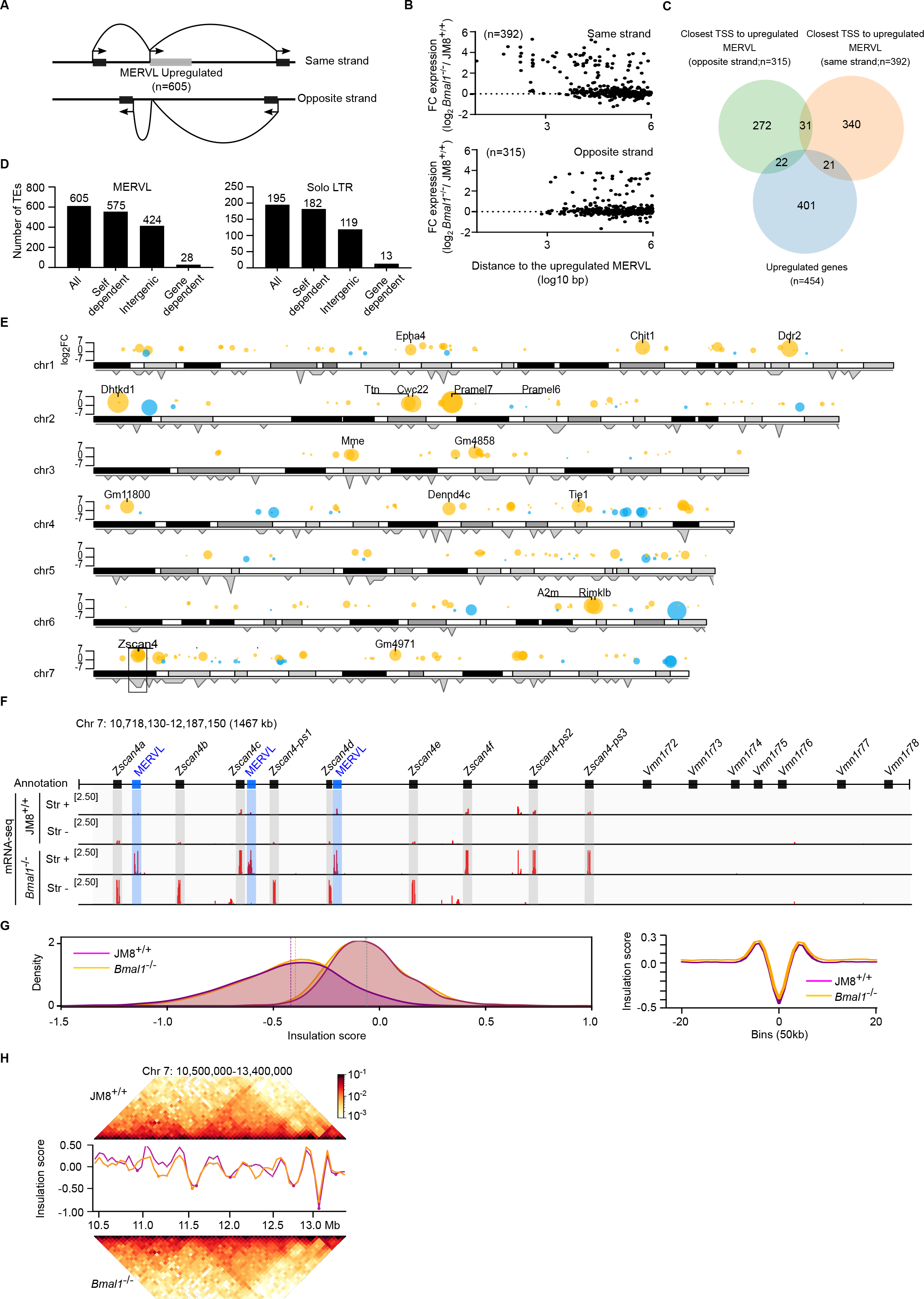
MERVL TEs and protein coding genes are transcriptionally induced at the Zscan4 locus (A) Scheme illustrating the approach to identify the closest annotated gene to transcriptionally induced MERVL elements in Bmal1^-/-^ mESCs in a strand-specific way. Starting from the genomic position of upregulated MERVLs (605 elements, FC>2, p<0.05) in Bmal1^-/-^ cells, the nearest annotated TSS upstream and downstream was identified and classified depending on the coding strand (same or opposite). (B) Plot showing the fold change expression (Bmal1^-/-^ vs JM8^+/+^) of genes identified in (A) relative to the distance to the upregulated MERVL copy. (C) Venn diagram showing the overlap between genes identified in (A) and protein coding genes transcriptionally induced in Bmal1^-/-^ cells (FC>2, p<0.05). (D) Classification of MERVL copies activated in Bmal1^-/-^ cells depending on whether they fall into an intergenic region or within protein coding genes that are transcriptionally induced in or not Bmal1^-/-^. Self-dependent are the sum of MERVL elements falling in intergenic and protein coding genes that are transcriptionally unaltered in Bmal1^-/-^. (E) Distribution of deregulated genes and MERVL elements across the indicated chromosomes. Circles indicate upregulated (yellow) or downregulated (blue) genes in Bmal1^-/-^ versus JM8^+/+^ cells. the statistical significance of the mis-regulation is indicated with the size of the circle. Grey histogram shows density of significantly upregulated MERVL copies. (F) Scheme showing annotated genes (black boxes) and complete MERVL elements (blue boxes) in the genomic interval chr7:10,706,720-12,139,219 (top panel). mRNA expression assayed by mRNA-seq in Bmal1^-/-^ and JM8^+/+^ mESCs in shown (bottom panel). (G) Plot comparing the distribution of the insulation score of strong boundaries or weak boundaries found in Bmal1^-/-^ cells in JM8^+/+^ and Bmal1^-/-^ cells (left panel). Plot comparing the average insulation score around strong boundaries identified in Bmal1^-/-^ cells (right panel). (H) Interaction maps of region containing Zscan4 genes in JM8^+/+^ and Bmal1^-/-^ cells and plots displaying the insulation score of the region in JM8^+/+^ (purple) and Bmal1^-/-^ (orange) cells. Dots mark strong boundaries.

**Figure S4.**
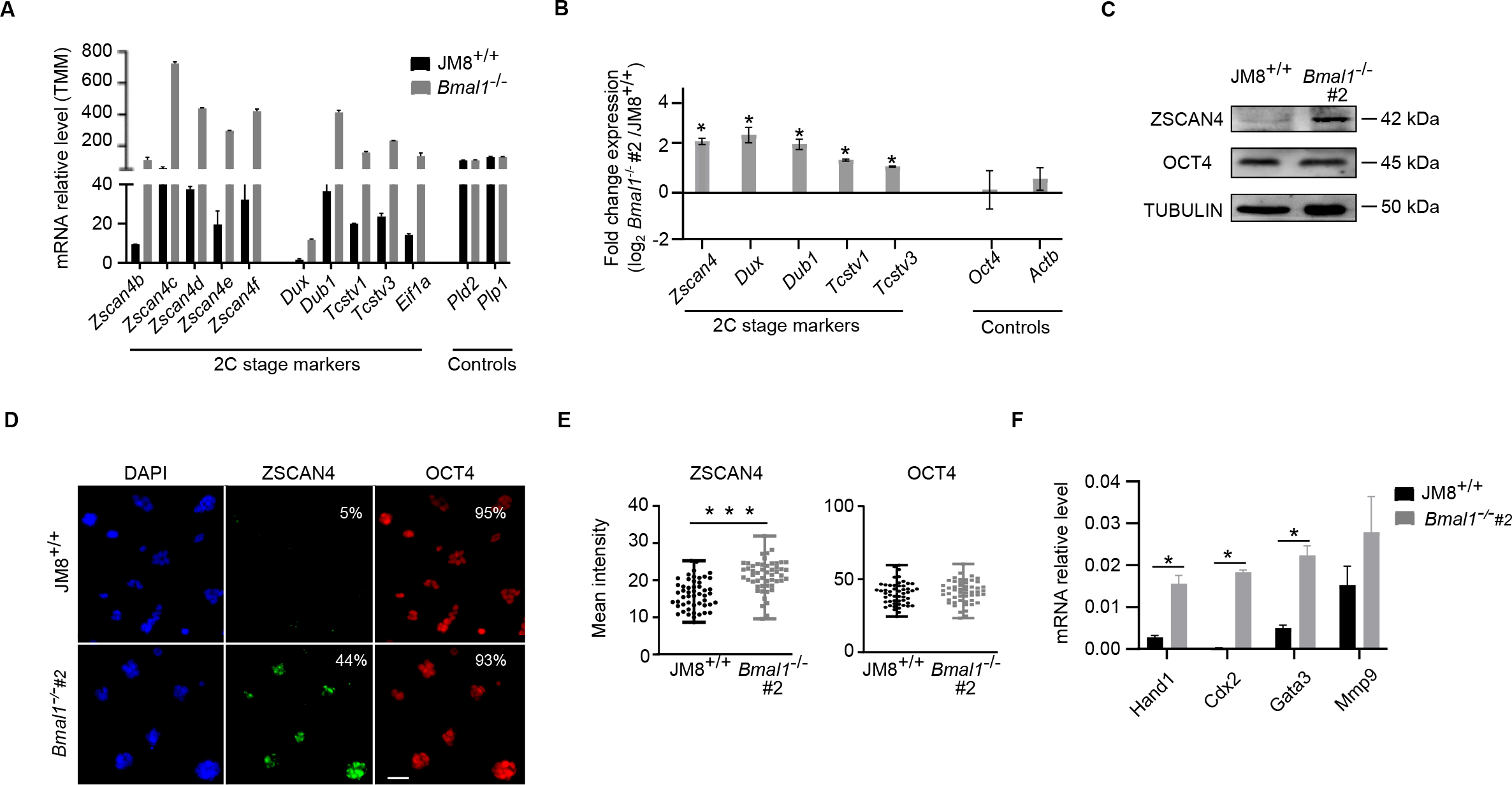
Bmal1^-/-^#2 mESCs acquire the expression of 2C-specific markers (A) Histogram showing the expression level of 2C-associated genes in JM8^+/+^ and Bmal1^-/-^ mESCs, as measured by mRNA-seq. Expression of Pld2 and Plp1 active genes is shown as a control. (B) Analysis of mRNA expression by RT-qPCR of 2C-associated genes in Bmal1^-/-^#2 relative to JM8^+/+^ mESCs. Gene expression was normalized to housekeeping genes (Hmbs, Yhwaz). Oct4 and Actb were included as controls. Mean ± SEM of three experiments is shown. Asterisks indicate p<0.05 in Mann-Whitney test. (C) Western Blot analysis of whole cells lysates measuring the expression of ZSCAN4 and OCT4 proteins in JM8^+/+^ and Bmal1^-/-^#2 mESCs. TUBULIN was used as a loading control. (D) Immunofluorescence images of DAPI, ZSCAN4 and OCT4 staining in Bmal1^-/-^#2 and JM8^+/+^ mESCs. The percentage of cells positive for ZSCAN4 or OCT4 labelling is indicated (n=52). Scale bar is 100 µm. (E) Plot showing the mean intensity of ZSCAN4 and OCT4 labelling in JM8^+/+^ (n=70 cells) and Bmal1^-/-^ #2 (n=70 cells) mESCs. Asterisks indicate p<0.0001 in Mann-Whitney test. (F) Gene expression analysis by RT-qPCR of trophectoderm-associated genes in JM8^+/+^ and Bmal1^-/-^ #2 mESCs. mRNA expression was normalized to Hmbs and Yhwaz housekeeping genes. Mean ± SEM of three experiments is shown. Asterisks indicate p<0.05 in Mann-Whitney test.

**Figure S5.**
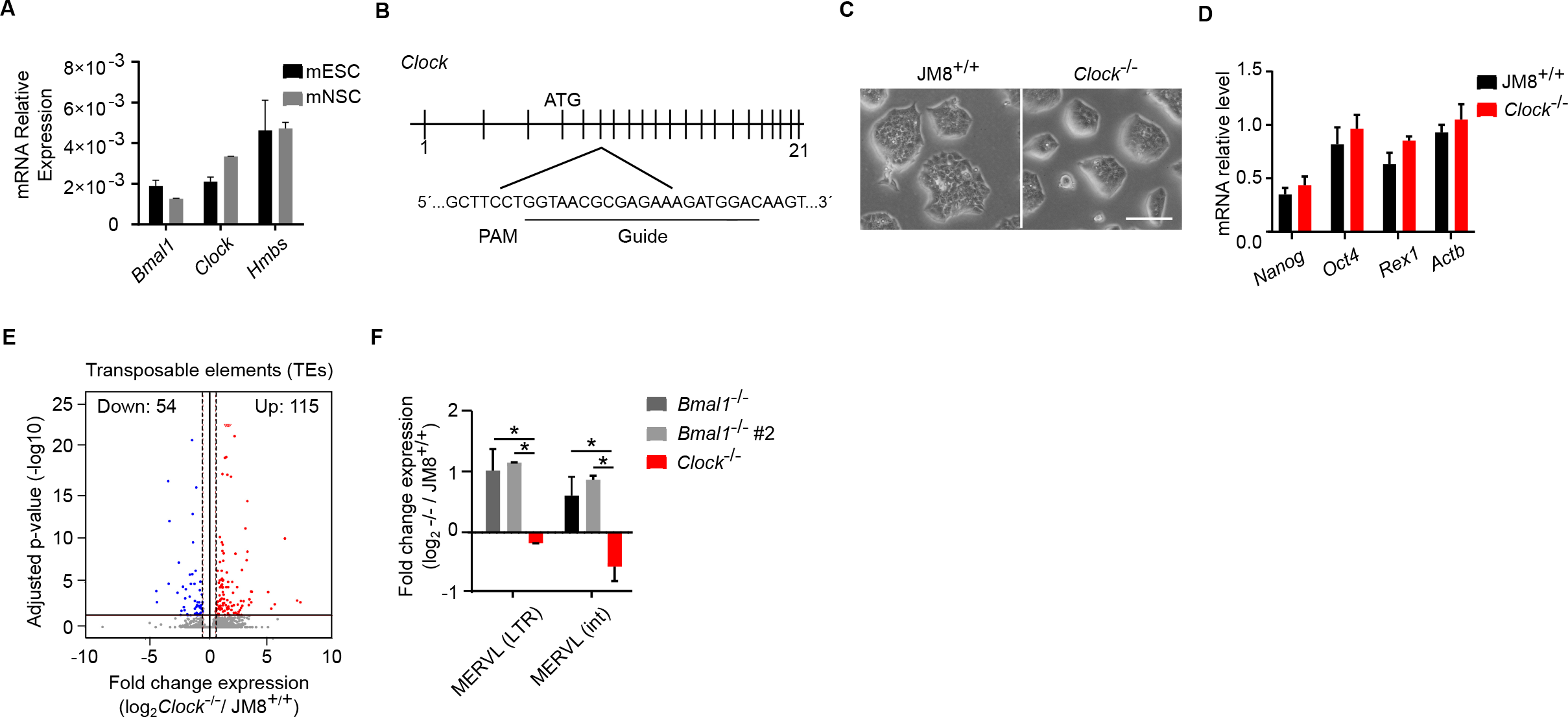
Clock-/- mESCs display unaltered colony morphology and expression of pluripotency markers (A) Expression analysis by RT-qPCR of Bmal1 and Clock genes in JM8 mESCs and in vitro-differentiated mNSCs. Hmbs provides a control. Expression was calculated relative to Gapdh and 18S genes. Mean ± SEM of three experiments is shown. (B) Scheme of the strategy used to generate a Clock^-/-^ mESC line using CRISPR/Cas9. Exon six of the Clock gene was targeted. The position of the ATG on exon 4, the DNA sequence targeted by the guide RNA, and the localization of the PAM sequence are shown. (C) Brightfield microscopy images of JM8^+/+^ and Clock^-/-^ mESCs colonies. Scale bar is 100 µm. (D) Expression analysis by RT-qPCR of pluripotency-associated genes in JM8^+/+^ and Clock^-/-^ mESCs. Expression was calculated relative to Hmbs and 18S genes. Mean ± SEM of three experiments is shown. Expression of Actb was used a control. (E) Volcano plot analysing the misexpression of TEs in Clock^-/-^ relative to parental JM8^+/+^ mESCs. TEs that are upregulated (FC>2, p-value<0.05) or downregulated (FC<2, p-value<0.05) are labelled in red and blue respectively, while non changing ones are labelled in grey. (F) RTq-PCR analysis of the expression of RNA from LTR or from the internal (int) region of MERVL elements in Bmal1^-/-^, Bmal1^-/-^#2, Clock^-/-^ mESCs. mRNA expression is normalized relative to the expression of Hmbs and Yhwaz genes and their matched parental cell lines. Mean ± SEM of three experiments is shown. Asterisks indicate *p<0.05 in Mann-Whitney test.

Table S1. List of genes and TEs upregulated in Bmal1-/- and Clock-/- mESCs.

Table S2. List of proteins identified by mass-spectrometry that interact with FLAG-BMAL1 in mESCs.

Table S3. List of reagents and datasets used in this study.

